# Loss of either RASSF1A alone or in combination with Caveolin-1 inhibition is associated with different premalignant histopathological alterations in the mammary glands of transgenic mice

**DOI:** 10.64898/2026.07.14.738049

**Authors:** Cristina L. Cotarelo, Hanna T. Weber, Sven Roßwag, Tabea Wagner, Ina Schäfer, Jonathan P. Sleeman, Sonja Thaler

## Abstract

Analyses of human breast carcinomas (BCs) and premalignant breast lesions show that the loss of RASSF1A is an early event in the development of ER+ BCs, which correlates linearly with malignant progression. This observation suggests that RASSF1A inhibition is important for the development and progression of ER+ BCs. In addition to RASSF1A, concurrent caveolin-1 (Cav-1) inhibition may further promote ER+ breast carcinogenesis. In the present study, transgenic *Rassf1a*-/- and *Cav-1*(-/-) single as well as *Rassf1a*-/-, *Cav-1*(-/-) double knockout mice were used to investigate the impact of single or combined *Rassf1a* and *Cav-1* inactivation on BC initiation. Loss of either one or both proteins led to different, pre-malignant histopathological alterations within the mammary glands of the mice, but not to fully developed BC, confirming that Rassf1a and Cav-1 are both important for maintaining the integrity of mammary gland epithelial structure, but suggesting that further intracellular changes or extracellular factors are required for the development of luminal BC when both genes are lost.

## Introduction

Estrogen receptors (ERs) exist as two different isoforms referred to as ERα and ERβ. They are steroid hormone receptors that act as ligand-activated transcription factors (Chen et al., 2022; Musgrove and Sutherland, 2009). Expression of the ERα is present in around 70-80% of all human breast carcinomas, whereas in normal breast tissue, ERα positive cells (hereafter referred to as ER+ cells) are in the minority (Shoker et al., 1999; Sørlie et al., 2001). The ERα is of central importance for the development of the mammary gland, and acts to regulate the proliferation and differentiation of normal breast tissue (Bocchinfuso et al., 2000). However, the ERα plays also an important role in the development of luminal breast cancer (Bocchinfuso and Korach, 1997; Hewitt et al., 2002; Shoker et al., 2000, 1999). Loss or functional inactivation of proteins controling ERα expression and function in the mammary gland is therefore an important and initial event during breast cancer development.

The ERα mediates estrogen-dependent effects on proliferation, differentiation, transformation and tumorigenesis (Bocchinfuso and Korach, 1997; Hewitt et al., 2002; Shoker et al., 2000, 1999) through so-called genomic and non-genomic activities (Chen et al., 2022; Musgrove and Sutherland, 2009; Siersbæk et al., 2018). In the nucleus, ERα undergoes conformational changes upon binding to estrogen that allow it to interact with co-activators and to bind to DNA, thereby promoting transcription of ERα-target genes directly through genomic mechanisms (Chen et al., 2022; Musgrove and Sutherland, 2009; Siersbæk et al., 2018). ERs localized to the plasma membrane also influence the activity of signalling pathways through non-genomic mechanisms (Chen et al., 2022; Musgrove and Sutherland, 2009; Siersbæk et al., 2018). For example, the ERα can be associated with the cell membrane through attachment to the scaffold protein caveolin-1 (Evinger and Levin, 2005; Razandi et al., 2002), and thereby forms complexes with G proteins, receptor tyrosine kinases (e.g. EGFR and IGFR-1) and non-receptor tyrosine kinases such as c-src (Migliaccio et al., 2002) and Ras (Migliaccio et al., 1996). This triggers signal transduction pathways that mainly evolve from G protein activation, leading to activation of downstream effectors such as mitogen activated protein kinase ERK (MAPK) (Migliaccio et al., 1996), phosphatidylinositol-3-OH kinase (PI3-K) and AKT (Lee et al., 2005; Sun et al., 2001). AKT in turn regulates multiple biological functions, including gene expression, cell cycle progression, survival, insulin-induced metabolic signals, endocytosis, cell transformation and oncogenesis (Manning and Toker, 2017).

Cell autonomous anti-tumor mechanisms such as apoptosis and senescence that protect cells from malignant transformation are executed by the tumor suppressor p53 (Lowe et al., 2004). The ERα can inhibit p53 activity through direct interaction with the protein (Liu et al., 2006), and indirectly through occupying binding sites within the promoter regions of p53 target genes (Bailey et al., 2012) or through inhibition of p53-mediated repression of genes (Sayeed et al., 2007). Some target genes of ERα encode proteins that can promote stem cell-like properties in breast cancer cells, such as FOXM1, as well as proteins that are capable of cooperative malignant transformation such as Bcl2 and c-Myc (Bergamaschi et al., 2014; Dubik and Shiu, 1992; Millour et al., 2010; Perillo et al., 2000; Planas-Silva et al., 2007; Strasser et al., 1990; Yang et al., 1999). Thus, ERα plays a central role in breast cancer initiation by suppressing mechanisms that counteract malignant transformation, and can also promote drug resistance, recurrence and metastasis (Bergamaschi et al., 2014; Dubik and Shiu, 1992; Millour et al., 2010; Perillo et al., 2000; Planas-Silva et al., 2007; Strasser et al., 1990; Yang et al., 1999). Dysregulated ERα function and activity therefore can cause pathological changes in breast tissue such as premalignant intraductal hyperplasia and the promotion of breast carcinogenesis (Chen et al., 2022; Couse and Korach, 1999).

The Ras association domain family 1 isoform A (RASSF1A) is a tumor suppressor whose inactivation is implicated in the development of many human cancers (Burbee et al., 2001; Dammann et al., 2001). The most common mechanism for loss or reduction of RASSF1A expression and function is transcriptional silencing through hypermethylation of a CpG island upstream of exon 1α in the *RASSF1* gene (Burbee et al., 2001; Dammann et al., 2001). DNA methylation analyses of breast carcinomas and pre-malignant breast lesions demonstrate that epigenetic alterations of the *RASSF1A* promoter occur preferentially in ER+ breast tumors (Cho et al., 2012; Feng et al., 2007; Grawenda and O’Neill, 2015; Kajabova et al., 2013; Shinozaki et al., 2005; Sunami et al., 2008; Xu et al., 2012), and that loss of RASSF1A is an early event in the course of breast tumorigenesis, which correlates linearly with malignant progression (DeVaux and Herschkowitz, 2018; Lehmann et al., 2002; Park et al., 2011; van Hoesel et al., 2013). These observations suggest that loss of RASSF1A is likely to be a key step in the development and/or progression of luminal ER+ breast tumors.

Experimental studies show that restoration of RASSF1A expression suppresses the estrogen/ERα-driven growth of luminal ER+ breast cancer cells (Roßwag et al., 2021, 2020; Thaler et al., 2012), providing a rational explanation for why hypermethylation of the *RASSF1A* promoter and loss of RASSF1A expression occurs preferentially in human ER+ breast carcinomas. These studies demonstrate that RASSF1A expression suppresses ER+ breast cancer growth through a complex network that encompasses multiple signaling pathways (Roßwag et al., 2021, 2020; Thaler et al., 2012), supporting the notion that loss of RASSF1A is important for the development and progression of luminal ER+ breast cancer.

The structural protein caveolin-1 (here after referred to Cav-1) also plays a role in the regulation of ERα expression and function (Sotgia et al., 2006). Dominant-negative mutations of Cav-1 are found exclusively in human ERα+ breast carcinomas (Lee et al., 2002), suggesting a particular role for Cav-1 in suppressing luminal breast carcinogenesis. This notion is further supported by genetically engineered *Cav-1* knockout (*Cav-1*(-/-)) mice, which show pathological alterations in their mammary glands (Li et al., 2006). *Cav-1* also plays an important role in the regulation of signaling pathways that can lead to tumorigenesis if permanently activated. One of these intracellular signaling pathways is the Jak2/STAT5 pathway, which is activated by prolactin during pregnancy (Tian et al., 2020; Xie et al., 2002). Pregnant *Cav-1*-deficient mice exhibit accelerated lobuloalveolar development, premature milk production (precocious lactation), and constitutive activation of JAK2/STAT5 and the Ras/MAPK pathways (Park et al., 2002). However, *Cav-1*(-/-) mice do not spontaneously develop breast tumors (Lee et al., 2002) and further environmental or genetic insults are needed to produce full-blown breast tumors (Williams and Lisanti, 2005; Zhang et al., 2005).

Previously published experimental *in vivo* studies reported that *Rassf1a-/-* mice develop significantly more tumors than wild-type control mice (Tommasi et al., 2005), and that *Rassf1a*-/- and *Rassf1a*+/- mice showed increased tumor proliferation and tumor size compared to their wild-type counterparts (Tommasi et al., 2005). However, in these *in vivo* studies, additional exogenous factors such as exposure to carcinogens (e.g. benzo(a)pyrene and urethane) were required to induce skin and lung tumors within the *Rassf1a*-/- and *Rassf1a*+/- mice (Tommasi et al., 2005), or the mice had to be further genetically modified with an activated oncogene such as K-Ras to develop lung carcinomas (Schmidt et al., 2018). Although data based on patient samples suggest that the loss of RASSF1A plays a role in the development of ER+ luminal breast cancer, no *in vivo* studies have yet been performed to functionally investigate this association.

Conceivably, RASSF1A might act in concert with Cav-1 to inhibit ERα expression and function, or to suppress signalling pathways that are important for breast epithelial cell transformation and malignant progression. This hypothesis is based on several observations that indicate a role for Cav-1 in suppressing ERα expression (Li et al., 2006; Williams and Lisanti, 2005; Zhang et al., 2005). For example, *Cav-1*-deficient mice exhibit increased ERα expression and premalignant alterations in mammary epithelia such as pronounced lobular development with numerous acini per terminal ductal lobular unit, hyperplasia of the mammary epithelial lining and extensive fibrosis in contrast to matched wildtype counterparts, which show normal lobular development (Li et al., 2006). Furthermore, RASSF1A inhibits ERα in part through the same signalling pathways as Cav-1 such as MAPK and AKT (Thaler et al., 2012), further supporting the notion that both proteins might cooperate to inhibit ERα expression and function, or to suppress transformation-fostering signalling pathways. Thus, following mammary epithelial cell hyperplasia caused by *Cav-1* insufficiency, loss of *RASSF1A* might be a critical event that leads to the onset of mammary carcinogenesis.

In this study, we investigated these issues by crossing transgenic *Rassf1a*-/- mice (Tommasi et al., 2005) with *Cav-1* (-/-)-deficient mice (Razani et al., 2001). Mammary glands from female mice were analyzed for changes in their structure and cellular composition compared to controls, and to determine whether pre-malignant lesions appear earlier and more often in the *Rassf1a*-/- ;*Cav-1* (-/-)-deficient mice. We observed that loss of either one or both proteins leads to different (pre-malignant) histopathological alterations within the mammary glands, but not to malignant transformation and cancer formation. These observations confirm that Rassf1a and Cav-1 are required to maintain the integrity of the mammary epithelial architecture and to restrain hormone-driven premalignant changes, but show that their combined loss alone is not sufficient for malignant transformation. Further intracellular changes such as the activation of oncogenes or extracellular factors seem to be additionally required for malignant progression and the development of full-blown luminal breast carcinomas.

## Results

### Mammary glands from *Rassf1a/caveolin-1* wild-type, single knockouts and double knockouts show differences in size and morphology

In order to study the impact of *Rassf1a* and *Cav-1* inactivation on luminal breast carcinogenesis, the inguinal mammary glands (#4 and #5) of wild-type (*Rassf1a+/+;Cav-1*(*+/+*)), *Rassf1a* single knockout (*Rassf1a-/-;Cav-1*(*+/+*)), *Cav-1* single knockout (*Rassf1a+/+;Cav-1*(*-/-*)) and *Rassf1a*/*Cav-1* double knockout (*Rassf1a-/-;Cav-1*(*-/-*)) were excised at various time points following birth and used for whole mount preparation. Mammary glands were collected from female mice with an age of between 4 to 12 months. Whole mount staining demonstrated differences in size and phenotype of the mammary glands taken from the different genotypes. Compared to ducts in wild-type tissue, mammary glands from *Rassf1a-/-;Cav-1*(*+/+*) mice contained enlarged ducts with spike-like protrusions (Figure 1 (A) and (B)). However no differences in the size of the mammary fat pads were observed when age-matched wild-type and *Rassf1a-/-;Cav-1*(*+/+*) were compared,. Mammary fat pads from mice with a *Cav-1*-deficiency (*Rassf1a+/+;Cav-1*(*-/-*) and *Rassf1a-/-;Cav-1*(-/-)) were smaller in comparison to those from *Cav-1* (+/+) mice, and exhibited enlarged duct lumina and acinar structures (Figure 1 (A)-(D) left and right panels). Genotype-specific changes in the appearance of the mammary glands were age independent. Based on the whole mounts, no significant differences in the size of the mammary fat pads and the appearance of the mammary glands could be observed between comparable *Rassf1a*+/+;*Cav-1*(-/-) and *Rassf1a*-/-;*Cav-1*(-/-) mice (Figure 1 (C) and (D)).

**Figure 1.**
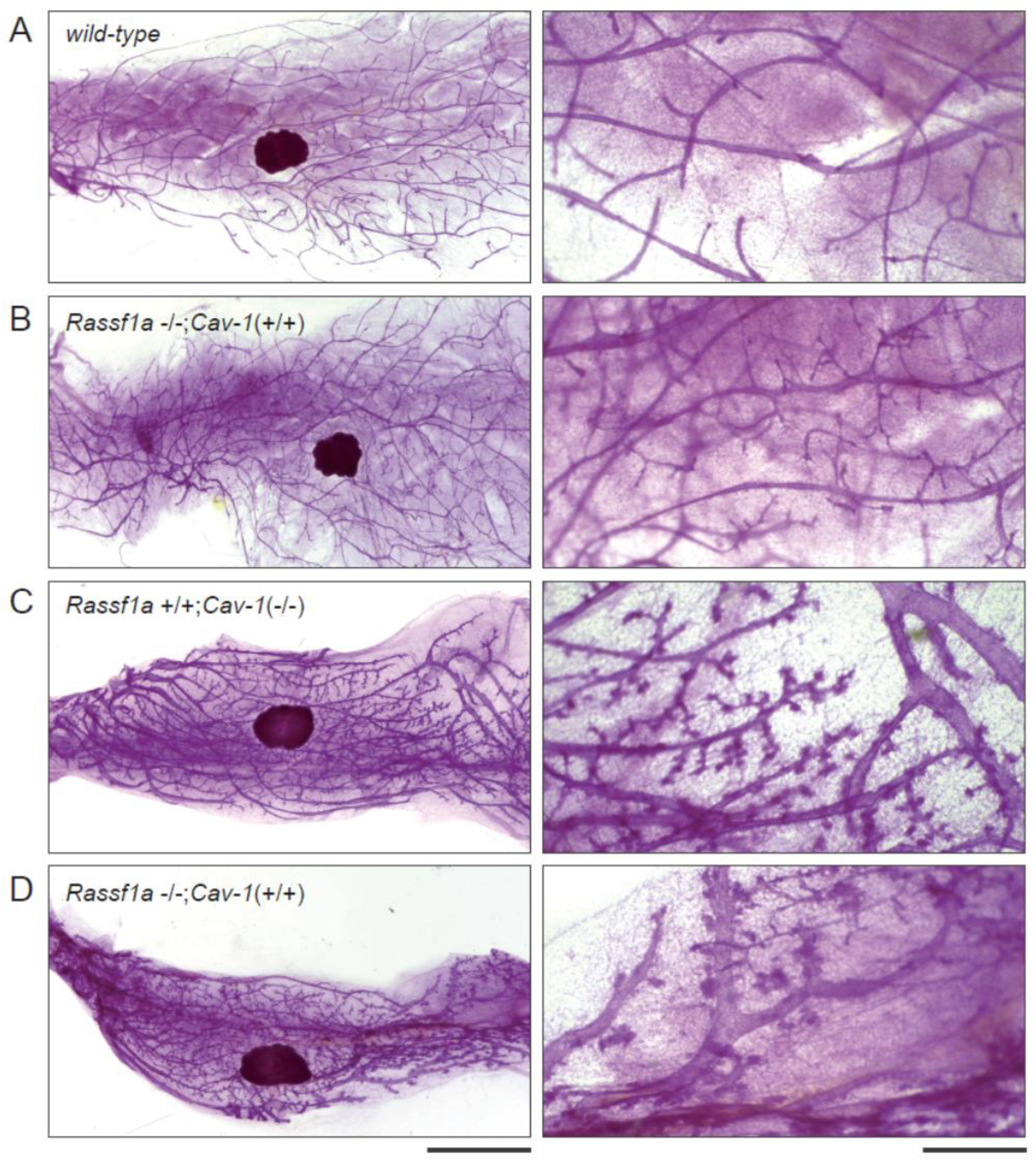
Mammary glands from *Rassf1a*/*Cav-1* wild-type, single knockouts and double knockouts show differences in size and appearances. Whole mount mammary glands were visualized by staining with carmine alum. (A)-(D) Representative pictures of stained mammary glands from comparable female mice of the same age. (A) *wild-type* (left and right panel) (B) *Rassf1a*-/-;*Cav-1*(+/+) (left and right panel) (C) *Rassf1a*+/+;*Cav-1*(-/-) (left and right panel) (D) *Rassf1a*-/-;*Cav-1*(-/-) (A),(B),(C),(D) left panel bar = 1 cm and right panel Bar = 0,5 µm

### Mammary glands from *Rassf1a/Cav-1* wild-type, single knockouts and double knockouts can be divided into different histopathological phenotypes

In order to study the consequences of *Rassf1a* and *Cav-1* inactivation for histopathological changes, inguinal mammary glands (#4 and #5) from comparable *Rassf1a+/+;Cav-1*(*+/+*), *Rassf1a-/-;Cav-1*(*+/+*), *Rassf1a+/+;Cav-1*(*-/-*) and *Rassf1a-/-;Cav-1*(*-/-*) female mice were again excised. To characterize changes in the histological structure and cellular composition of the mammary glands compared to appropriate controls, and to determine whether pre-malignant lesions and/or malignant lesions appear earlier and more often in the *Rassf1a*-/- and *Rassf1a*-/-;*Cav-1*(-/-)-deficient mice, the mammary glands from all genotypes were fixed in formalin and embedded in paraffin. After preparation of paraffin sections, tissue sections were H&E stained and histo-pathologically analyzed. Sections from formalin-fixed paraffin-embedded mammary glands taken from a total of 161 mice were analyzed (Table 1).

**Table 1.** Histological characteristics of mammary glands with different genotypes.

|  |  | Genotype |  |  |  |
| --- | --- | --- | --- | --- | --- |
| Histological Type | Cases (n=161) | *Rassf1a/Cav-1*<br>*+/+,+/+*<br>(n=56) | *Rassf1a/Cav-1*<br>*-/-,+/+*<br>(n=38) | *Rassf1a/Cav-1*<br>*+/+,-/-*<br>(n=38) | *Rassf1a/Cav-1*<br>*-/-,-/-*<br>(n=57) |
| Normal (N) | 33 | 23<br>(41,07%) | 10<br>(26,32%) | 0 | 0 |
| (N/DCH) | 39 | 33<br>(58,93%) | 4<br>(10,53%) | 2<br>(5,26%) | 0 |
| Ductal cystic hyperplastic (DCH) | 46 | 0 | 21<br>(55,26%) | 6<br>(15,79%) | 19<br>(33,33%) |
| (DCH/DCAH) | 5 | 0 | 1<br>(2,63%) | 1<br>(2,63%) | 3<br>(5,27%) |
| Ductal cystic acinaric hyperplastic (DCAH) | 64 | 0 | 2<br>(5,26%) | 29<br>(76,32%) | 35<br>(61,4%) |

Based on the histopathological changes found in the mammary glands of the different genotypes, three different histological phenotypes can be defined, and hereafter referred to as normal (N), ductal cystic hyperplastic (DCH) and ductal cystic acinaric hyperplastic (DCAH) (Figure 2 (A)-(C)).

**Figure 2.**
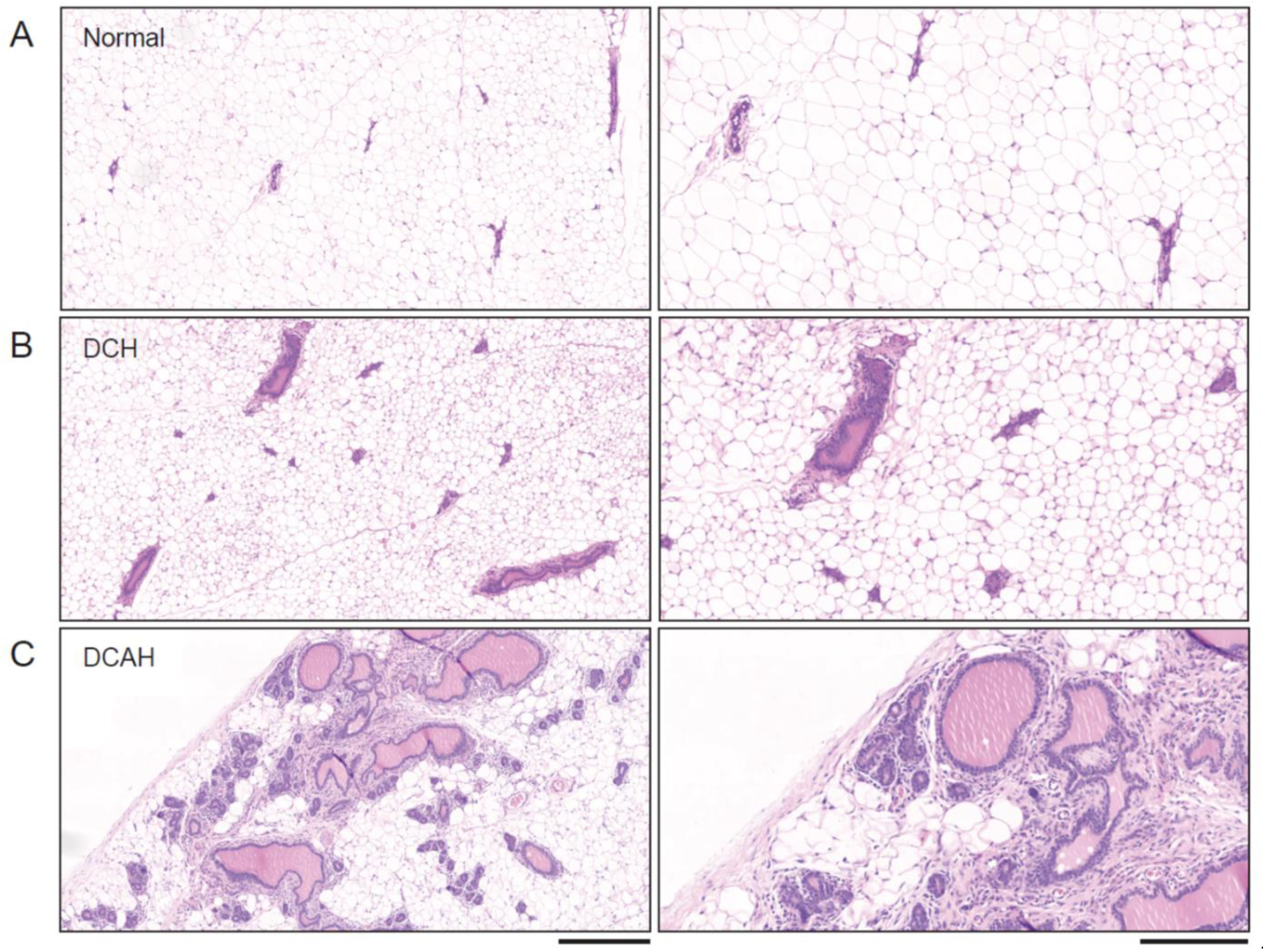
Histopathological phenotypes of the analyzed mammary glands. Representative pictures of H&E stained sections from paraffin-embedded formalin-fixed mammary tissue showing three different histopathological phenotypes that can be found in the mammary glands from the wild-type and *Rassf1a* and/or *Cav-1*-targeted mice (A) normal type (left and right panel), (B) ductal-cystic hyperplastic type (DCH) (left and right panel), (C) ductal-cystic and acinaric hyperplastic type (DCAH) (A)-(C) left panel Bar = 400µm; right panel Bar = 200µm

The N phenotype corresponds to normal ductal structures in the mammary gland without the formation of lobuloalveolar structures that normally only develop during pregnancy. The ducts have normal-sized lumens and are lined by a normal epithelial cell layer without any signs of epithelial hyperplasia. The DCH phenotype shows alterations in the mammary epithelium, such as over-sized ductal structures, cyst-like lumens and signs of epithelial hyperplasia. The DCAH phenotype is characterized by pronounced lobular development, enlarged ducts with over-sized, cyst-like lumens and with numerous acini per terminal ductal lobular unit, hyperplasia of the mammary epithelial lining and extensive fibrosis. The DCAH phenotype is consistent with alterations in the mammary glands of Cav-1-deficient mice reported previously (Li et al., 2006). In some mice, mammary glands contained transitional forms in which structural changes associated with two different histological phenotypes could be found (N/DCH and DCH/DCAH).

In the present study, 21 of the 38 *Rassf1a*-/-;*Cav-1*(-/-) mice that were analysed had DCH lesions, whereas no DCH lesions were found in the mammary glands of 56 wild-type mice. Therefore, *Rassf1a*-deficiency correlates significantly with the development of DCH (*p*<0.00001). In both the *wild-type* and the *Rassf1a*-/-;*Cav-1*(+/+) group, 33 mice exhibited normal, small, and slightly enlarged cystic ducts (N/DCH) with no significant differences between the groups (*p*=0.0706) (Table 1 and Table 2).

**Table 2.** Statistically significant differences exist in the appearance of ductal cystic hyperplasia (DCH) between the mammary glands from *Rassf1a*/*Cav-1* wild-type (+/+,+/+) and *Rassf1a* knockout *Cav-1* wild-type (-/-,+/+) mice.

| Genotype | Histological Type | Cases (n=94) | A | B | C |
| --- | --- | --- | --- | --- | --- |
| **Rassf1a/Cav-1**<br>**+/+,+/+**<br>**(n=56)** | Normal (N) | 23<br>(41,07%) | 23<br>(41,07%) | 56<br>(100%) | 23<br>(100%) |
|  | (N/DCH) | 33<br>(58,93%) | 33<br>(58,93%) |  |  |
|  | Ductal cystic hyperplastic (DCH) | 0 |  | 0 | 0 |
| **Rassf1a/Cav-1**<br>**-/-,+/+**<br>**(n=35)** | Normal (N) | 10<br>(28,57%) | 10<br>(71,43%) | 14<br>(40%) | 10<br>(32,26%) |
|  | (N/DCH) | 4<br>(11,43%) | 4<br>(28,57%) |  |  |
|  | Ductal cystic hyperplastic (DCH) | 21<br>(60%) |  | 21<br>(60%) | 21<br>(67,74%) |
| **Statistical significance** |  |  | p=0.0706<br>(n.s.) | p<0.00001 | p<0.00001 |

In 3 out of 38 *Rassf1a*-/-;*Cav-1*(+/+) mammary glands, either DCH/DCAH or DCAH was observed, whereas no wild-type mice had DCH/DCAH or DCAH within their mammary glands (Table 1). In the two *Cav-1*-deficient groups, the majority of mice displayed DCAH in their mammary glands (76.32% of mice in the *Rassf1a*+/+;*Cav-1*(-/-) and 61.4% in the *Rassf1a*-/-;*Cav-1*(-/-) group), whereas no mice with a normal phenotype were observed (Table 1). Significant differences were found in the appearance of DCH/DCAH and DCAH between mammary glands from *Rassf1a*-/-;*Cav-1*(+/+) and *Rassf1a*+/+;*Cav-1*(-/-) mice (*p*<0.00001). However, no significant differences in the presence of DCH in mammary glands from *Rassf1a*-/-;*Cav-1*(+/+) and *Cav-1*-deficient mice (*p*=0.6882) were observed (Table 3). Thus, *Cav-1*-deficiency significantly correlates with the development of DCAH. No statistically significant differences were found in the presence of DCH or DCAH when the mammary glands from *Rassf1a*+/+;*Cav-1*(-/-) and *Rassf1a*-/-;*Cav-1*(-/-) mice were compared (Table 4). Despite the fact that loss of either one or both tumor suppressors caused different (pre-malignant) histopathological alterations within the mammary glands, no cancer formation could be observed. This result suggests that additional events such as activation of oncogenes or external non-cell autonomous factors are needed in order for full-blown breast carcinomas to develop.

**Table 3.**
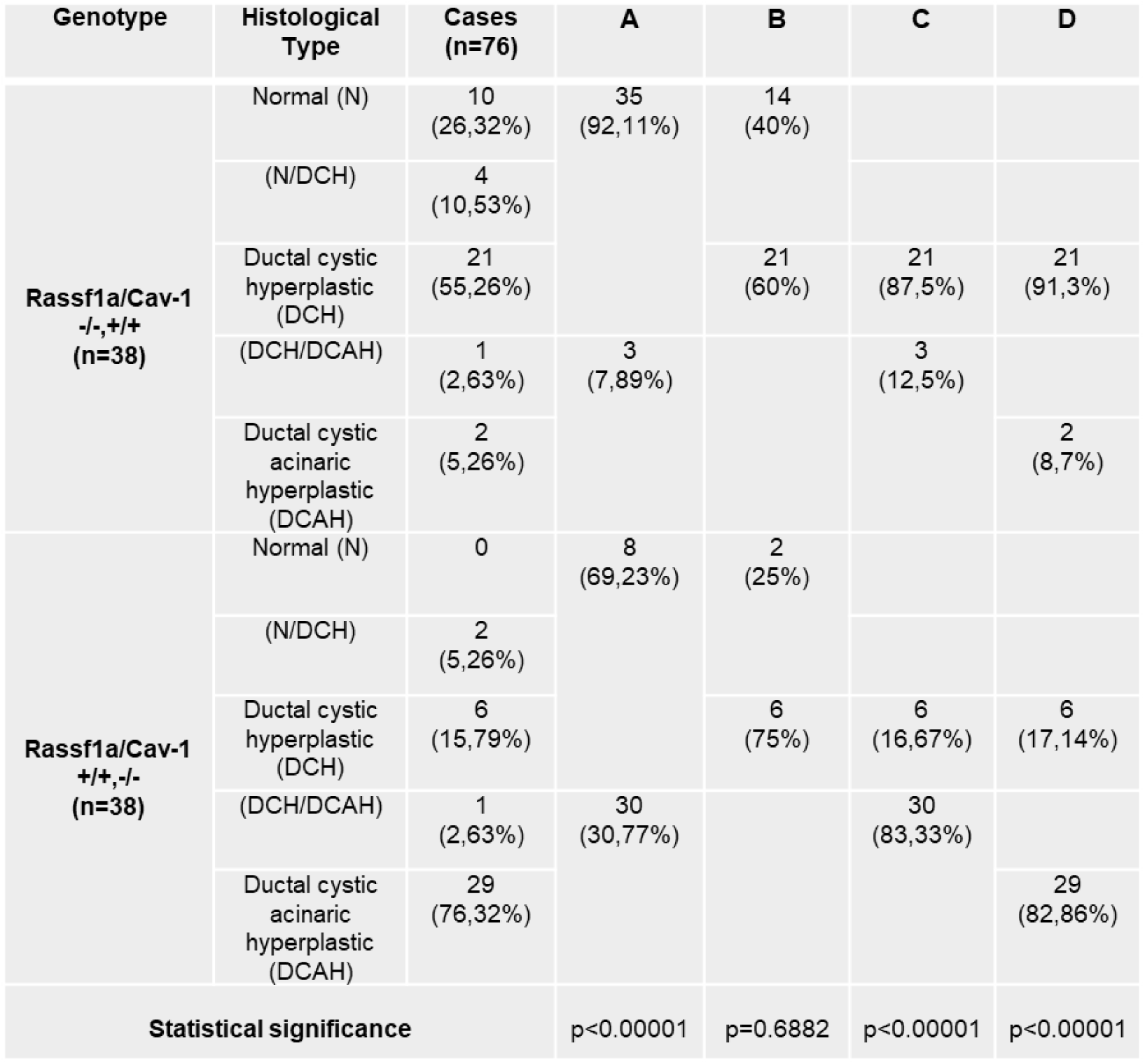
Statistically significant differences exist in the appearance of DCHA between the mammary glands from *Rassf1a*-/-;*Cav-1*(+/+) and *Rassf1a*+/+;*Cav-1*(-/-) mice.

**Table 4.**
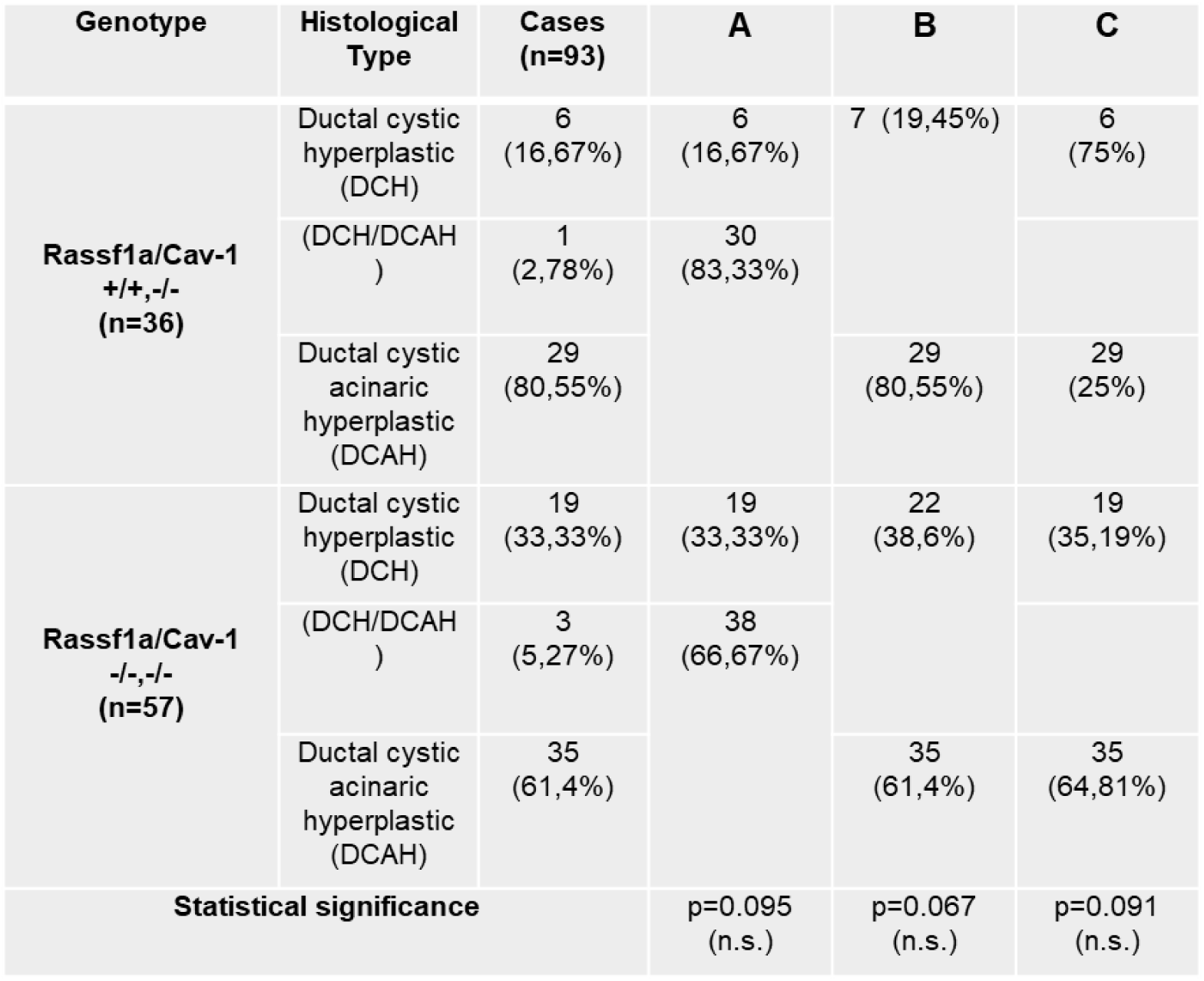
No statistically significant differences exist in the appearance of DCH and DCAH between mammary glands from *Rassf1a*+/+;*Cav-1*(-/-) and *Rassf1a*-/-;*Cav-1*(-/-) mice.

Isolated milk drops were found in the ducts and ductal-alveolar structures of mammary glands from *Rassf1a*-/-;*Cav-1*(+/+) mice, which was particularly pronounced in *Cav-1*-deficient *Rassf1a+/+*;*Cav-1(-/-)* and *Rassf1a*-/-;*Cav-1*(-/-) mice, indicating non-specific activation of the JAK2/STAT5 signaling pathway. This signaling pathway is normally only activated during pregnancy under the influence of prolactin (Tian et al., 2020; Xie et al., 2002). This observation is consistent with previous studies showing that Cav-1 loss leads to non-specific activation of the JAK2/STAT5 pathway (Park et al., 2002). The fact that some *Rassf1a*-/-;*Cav-1*(+/+) mice also had milk droplets in their mammary glands could potentially indicate that the loss of *Rassf1a* also influences the activation of the JAK2/STAT5 pathway.

Two *Rassf1a-/-*;*Cav-1*(-/-) mice, which were not listed in Table 1, exhibited massive cystic structures that were completely filled with milk drops (Figure 3 (A) and (B)). These females were neither pregnant nor lactating. Such a phenotype was only observed in the *Rassf1a-/-*;*Cav-1*(-/-) mice. This observation further supports the notion that loss of both Cav-1 and Rassf1a fosters milk production in the mammary gland.

**Figure 3.**
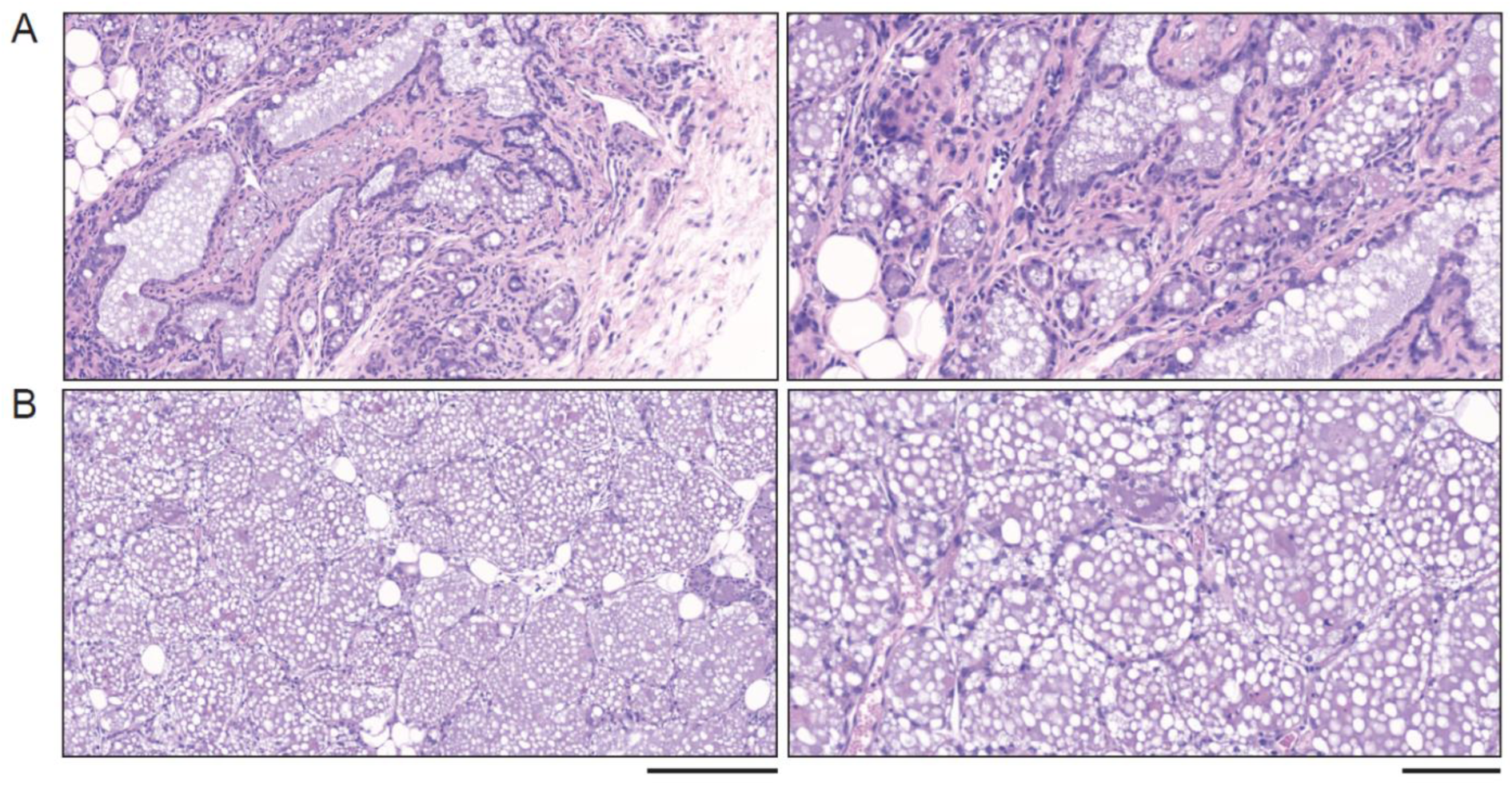
Secretory phenotype observed in mammary glands of *Rassf1a-/-/Cav-1(-/-)* mice. (A)+(B) Representative pictures of H&E-stained sections from paraffin-embedded formalin-fixed mammary tissue from two non-pregnant, non-lactating *Rassf1a*-/-/*Cav-1*(-/-) mice showing an extensive secretory phenotype. (A) and (B) left panel Bar = 200 µm; (A)+(B) right panel Bar = 100µm

### Mammary glands from pregnant *Rassf1a/Cav-1* wild-type, single knockouts and double knockouts show no signs of malignancy

To test the possibility that concomitant loss of *Rassf1a* and *Cav-1* might be a critical event that leads to the onset of malignant transformation and mammary carcinogenesis during pregnancy, mammary glands from pregnant mice were excised at either between day 13,5 and day 14 or between day 17,5 and day 18. To characterize changes in the histological structure and cellular composition compared to appropriate controls, and to determine whether pre-malignant lesions and/or malignant lesions appear in the *Rassf1a*-/- and *Rassf1a*-/-;*Cav-1*(-/-)-deficient mice, the mammary glands from all genotypes were fixed in formalin and embedded in paraffin. After preparation of paraffin sections, tissue sections were H&E stained and histopathologically analyzed. Sections from formalin-fixed paraffin-embedded mammary glands taken from a total of 23 pregnant mice were analyzed (Table 5).

**Table 5.** Mammary glands from pregnant mice with different genotypes.

|  |  | Genotype |  |  |  |
| --- | --- | --- | --- | --- | --- |
| day of pregnancy | Cases (n=23) | *Rassf1a/Cav-1*<br>+/+,+/+<br>(n=8) | *Rassf1a/Cav-1*<br>-/-,+/+<br>(n=5) | *Rassf1a/Cav-1*<br>+/+,-/-<br>(n=5) | *Rassf1a/Cav-1*<br>-/-,-/-<br>(n=5) |
| **13.5 - 14** | 9 | 3 | 2 | 2 | 2 |
| **17.5 - 18** | 14 | 5 | 3 | 3 | 3 |

The alveolar switch in pregnant mice is a hormonally driven genetic program, which triggers rapid remodeling of the ductal epithelium into milk-secreting lobuloalveolar units (Oakes et al., 2006). It coordinates proliferation, differentiation and structural changes, which are necessary for lactation. The hormones progesterone and prolactin initiate this developmental switch, fostering the expansion of secretory alveoli (Brisken et al., 2002). Histopathological analysis of the H&E-stained paraffin sections showed typical morphological changes in the mammary glands during pregnancy due to the alveolar switch in all different genotypes. Characteristic changes such as the development of milk-secreting lobuloalveolar units could be observed in all genotypes (Figure 4 (A)-(D)).

**Figure 4.**
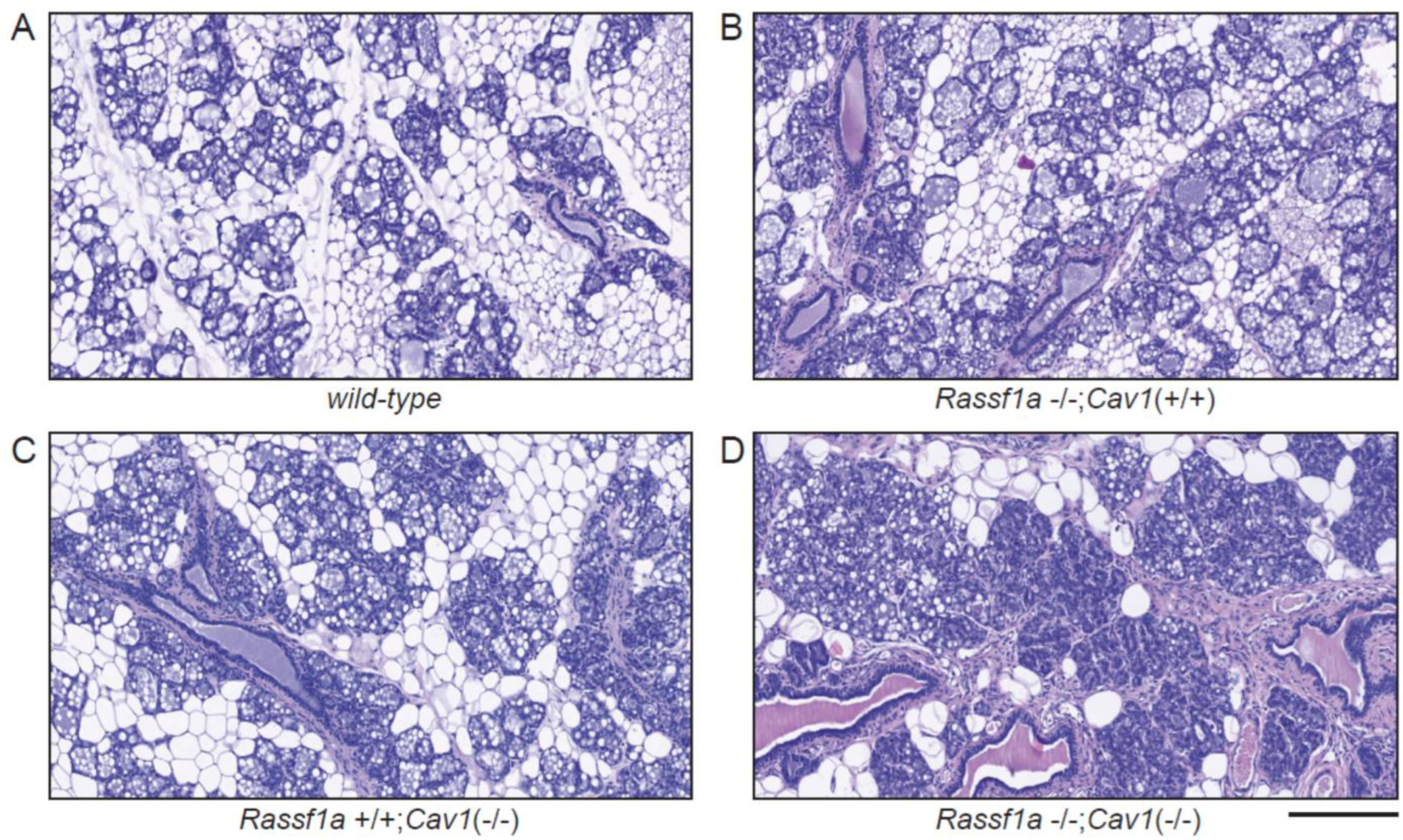
Histopathological analysis of the mammary glands from pregnant mice. Representative pictures of H&E-stained sections from paraffin-embedded formalin-fixed mammary tissue taken from pregnant wildtype and *Rassf1a* and/or *Cav-1*-targeted mice between day 17.5 and day 18 (A) wildtype with typical lobuloalveolar units (normal appearance), (B) *Rassf1a*-/-;*Cav-1*(+/+) with lobuloalveolar units and DCH, (C) *Rassf1a*+/+;*Cav-1*(-/-) and (D) *Rassf1a*-/-;*Cav-1*(-/-) (C) and (D) both *Cav-1*-deficient genotypes have lobuloalveolar units, pronounced lobular development, enlarged ducts with over-sized, cyst-like lumens, hyperplasia of the mammary epithelial lining and extensive fibrosis. Bar = 200 µm

Nevertheless, the histopathological alterations that were observed in the mammary glands of non-pregnant *Rassf1a*-/-;*Cav-1*(+/+), *Rassf1a*+/+;*Cav-1*(-/-) and *Rassf1a*-/-;*Cav-1*(-/-) mice such as enlarged, over-sized ductal lumens with epithelial hyperplasia and fibrosis were also evident in these genotypes during pregnancy (Figure 2 and Figure 3). Although *Cav-1* plays an important role in regulating JAK2/STAT5 and Ras/MAPK signalling that can contribute to tumorigenesis, *Cav-1*-deficient mice do not develop breast cancer (Park et al., 2002). In the study presented here, neither *Rassf1a*+/+;*Cav-1*(-/-) nor *Rassf1a*-/-;*Cav-1*(-/-) mice developed breast cancer during pregnancy. These results demonstrate that concomitant loss of Rassf1a is not sufficient to initiate breast cancer development under the influence of hormones such as prolactin and progesterone.

### Immunohistological staining of mammary glands from *Rassf1a/Cav-1* wild-type, single knockouts and double knockouts show differences in the expression of ERα

*Cav-1*-deficient mice exhibit increased ERα expression in mammary epithelial cells, and mammary glands from *Cav-1* (-/-) mice are functionally hypersensitive to the effects of estrogen exposure (Mercier et al., 2009). RASSF1A inhibits ERα expression and function in parts through the same signalling pathways as Cav-1, such as MAPK and AKT (Thaler et al., 2012). To investigate the hypothesis that the Rassf1a and Cav-1 proteins might act in concert to inhibit ERα expression, inguinal mammary glands (#4 and #5) from *Rassf1a+/+;Cav-1*(*+/+*), *Rassf1a-/-;Cav-1*(*+/+*), *Rassf1a+/+;Cav-1*(*-/-*) and *Rassf1a-/-;Cav-1*(*-/-*) female mice were excised, fixed and sectioned. ERα expression in the different genotypes was then quantified following immunohistochemical staining with an antibody against mouse ERα. Sections from formalin-fixed paraffin-embedded mammary glands taken from a total of 44 mice with an age of between 4 and 12 months were stained and analyzed (Table 6).

**Table 6.**
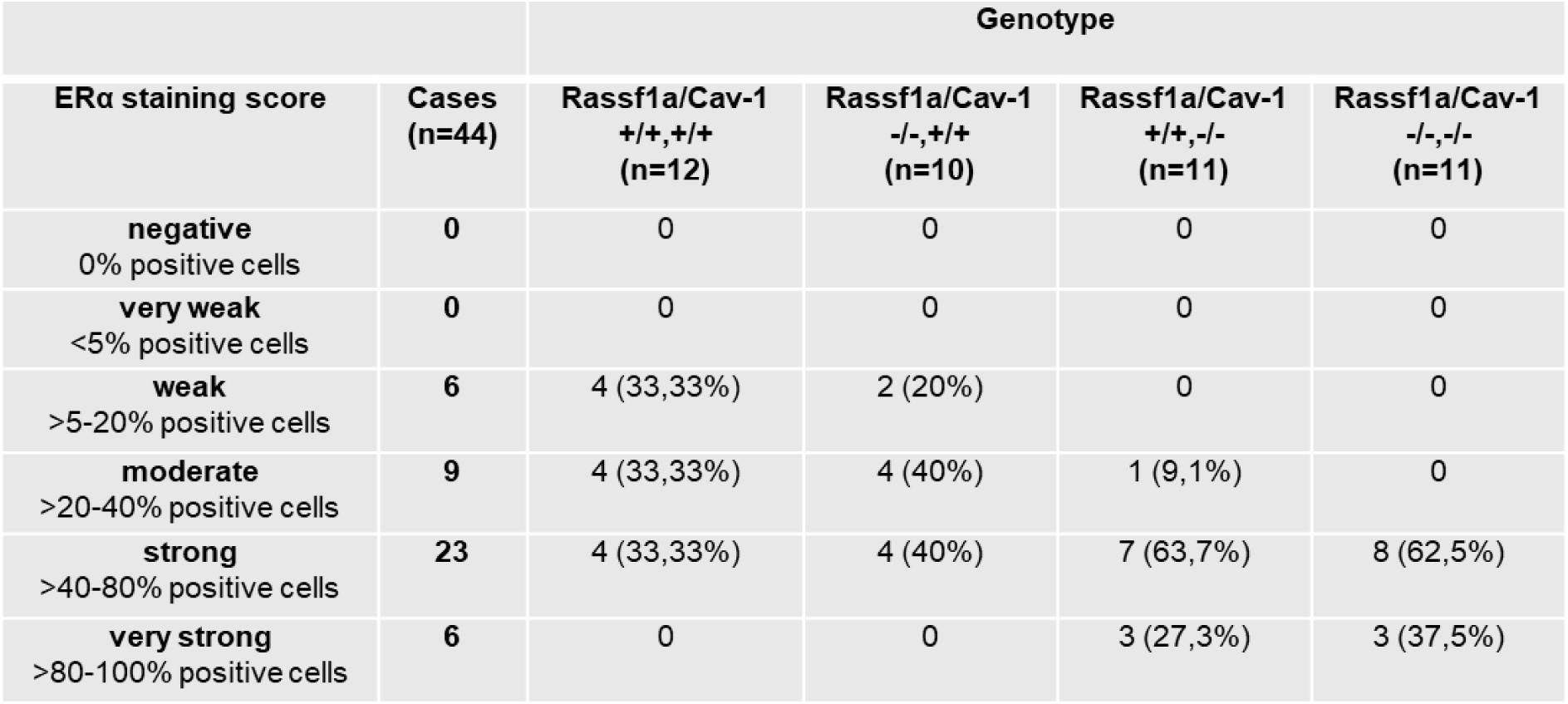
ERα expression in mammary glands from mice with different genotypes.

|  |  | Genotype |  |  |  |
| --- | --- | --- | --- | --- | --- |
| ERα staining score | Cases (n=44) | *Rassf1a/Cav-1*<br><sup>+/+</sup>,<sup>+/+</sup><br>(n=12) | *Rassf1a/Cav-1*<br><sup>-/-</sup>,<sup>+/+</sup><br>(n=10) | *Rassf1a/Cav-1*<br><sup>+/+</sup>,<sup>-/-</sup><br>(n=11) | *Rassf1a/Cav-1*<br><sup>-/-</sup>,<sup>-/-</sup><br>(n=11) |
| **negative**<br>0% positive cells | 0 | 0 | 0 | 0 | 0 |
| **very weak**<br><5% positive cells | 0 | 0 | 0 | 0 | 0 |
| **weak**<br>>5-20% positive cells | 6 | 4 (33,33%) | 2 (20%) | 0 | 0 |
| **moderate**<br>>20-40% positive cells | 9 | 4 (33,33%) | 4 (40%) | 1 (9,1%) | 0 |
| **strong**<br>>40-80% positive cells | 23 | 4 (33,33%) | 4 (40%) | 7 (63,7%) | 8 (62,5%) |
| **very strong**<br>>80-100% positive cells | 6 | 0 | 0 | 3 (27,3%) | 3 (37,5%) |

Representative pictures demonstrating the expression of ERα in mammary epithelia cells in the different mice genotypes are shown in Figure 5 (A)-(D). Tissue sections from a human ER- and a human ER+ mammary carcinoma were used as negative and positive controls on each slide. Analysis of the IHC-stained mammary glands used a scoring system in which the percentage of ERα-positive epithelial cells in the ducts were evaluated as described in Table 6. The distribution patterns of ERα-expressing cells in the mammary glands did not differ significantly between wild-type and *Rassf1a*-/-;*Cav-1*(+/+) mice. In addition, weak, moderate and strong staining was equivalently distributed in both genotypes (Table 6). The number of ERα-expressing cells in the mammary glands of the *Cav-1*-deficient *Rassf1a+/+;Cav-1*(*-/-*) and *Rassf1a-/-;Cav-1*(*-/-*) mice did significantly differ between wild-type and *Rassf1a*-/-;*Cav-1*(+/+) mice (Table 6). However, the distribution patterns of ERα-expressing cells in the mammary glands did not differ significantly between *Rassf1a+/+;Cav-1*(*-/-*) and *Rassf1a-/-;Cav-1*(*-/-*) mice. These results confirm that *Cav-1*-deficiency leads to increased ERα expression, as described previously (Mercier et al., 2009; Sotgia et al., 2006). With the exception of one mouse, *Cav-1*-deficient *Rassf1a+/+;Cav-1*(*-/-*) and *Rassf1a-/-;Cav-1*(*-/-*) mice have mammary glands that either contain more than 40% ERα-expressing epithelial cells (strong) or mammary glands that have 80%-100% ERα-expressing epithelial cells (very strong). These results should, however, be interpreted cautiously as the phase of the estrus cycle of the *Cav-1*-deficient mice at the time of mammary gland removal was not determined, which may account at least in part for these observations.

**Figure 5.**
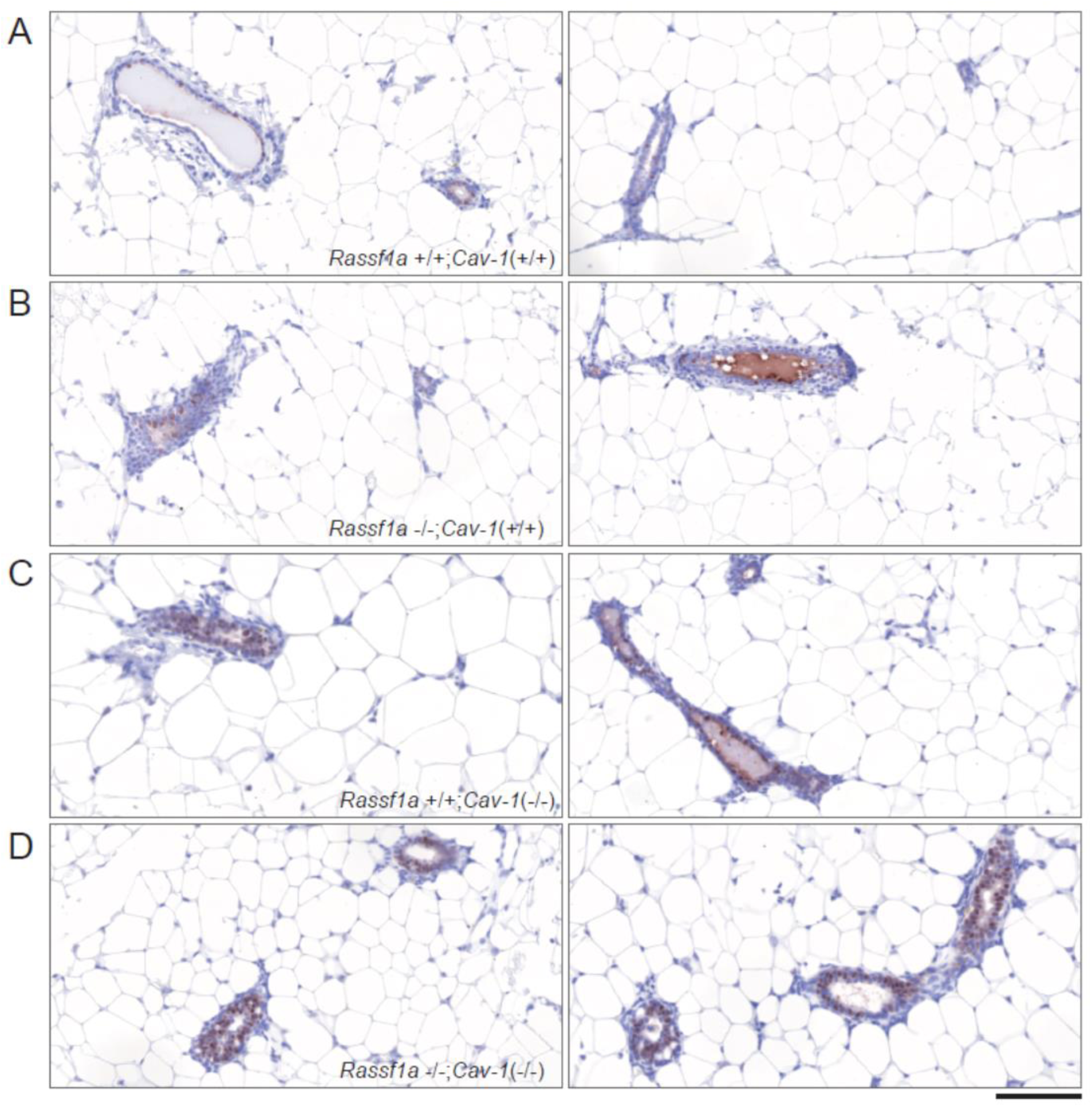
Immunohistochemical staining to analyse ERα expression in the mammary glands from mice with different phenotypes. Representative pictures of immunohistochemically-stained sections from paraffin-embedded formalin-fixed mammary tissue from wildtype and *Rassf1a* and/or *Cav-1*-targeted mice. (A) *wild-type*, (B) *Rassf1a*-/-;*Cav-1*(+/+), (C) *Rassf1a*+/+;*Cav-1*(-/-) and (D) *Rassf1a*-/-;*Cav-1*(-/-). (A) – (D) left and right Bar = 200 µm

The ERα is significantly down-regulated in the mammary gland epithelium during pregnancy in mice, and ERα expression decreases to a very low level, reaching a minimum around day 14 of gestation (Saji et al., 2000). To investigate whether *Cav-1*-deficiency and/or *Rassf1a*-deficiency plays a role in the downregulation of ERα during pregnancy, mammary glands were isolated from pregnant mice of all genotypes and ERα expression was analyzed. Mammary glands from 16 mice in total with an age between day 14 to 18 days were analysed (Table 7).

**Table 7.** Formation of ductal structures in the fat pads of male mice with different genotypes.

|  |  | Genotype |  |  |  |
| --- | --- | --- | --- | --- | --- |
| ERα staining score | Cases (n=16) | Rassf1a/Cav-1 +/+,+/+ (n=6) | Rassf1a/Cav-1 -/-,+/+ (n=4) | Rassf1a/Cav-1 +/+, -/- (n=3) | Rassf1a/Cav-1 -/-, -/- (n=3) |
| **negative**<br>0% positive cells | 7 | 2 (33,3%) | 2 (50%) | 1 (33,3%) | 2 (66,7%) |
| **very weak**<br><5% positive cells | 9 | 4 (66,7%) | 2 (50%) | 2 (66,7%) | 1 (33,3%) |
| **weak**<br>>5-20% positive cells | 0 | 0 | 0 | 0 | 0 |
| **moderate**<br>>20-40% positive cells | 0 | 0 | 0 | 0 | 0 |
| **strong**<br>>40-80% positive cells | 0 | 0 | 0 | 0 | 0 |
| **very strong**<br>>80-100% positive cells | 0 | 0 | 0 | 0 | 0 |

| Male mammary fat pads | Cases<br>(n=21) | *Rassf1a*/*Cav-1*<br>+/+,+/+ (n=5) | *Rassf1a*/*Cav-1*<br>-/-,+/+ (n=5) | *Rassf1a*/*Cav-1*<br>+/+,-/- (n=5) | *Rassf1a*/*Cav-1*<br>-/-,-/- (n=6) |
| --- | --- | --- | --- | --- | --- |
| no mammary gland formation | 10 | 5 (100%) | 1 (20%) | 4 (80%) | 0 |
| Mammary gland formation | 11 | 0 | 4 (80%) | 1 (20%) | 6 (100%) |
| Mammary gland ≥ 1 cm | 3 | 0 | 0 | 0 | 3 (50%) |

No differences in the number of ERα-expressing cells in the mammary glands between the different genotypes were observed. This result demonstrates that mechanisms leading to down-regulation of ERα during pregnancy are functional even in the absence of *Cav-1* or in the concomitant absence of *Cav-1* and *Rassf1a* (Table 7).

### Mammary glands from male *Rassf1a/Cav-1* wild-type, single knockouts and double knockouts show differences in duct formation and elongation

The mammary glands of females from *Rassf1a*-/-;*Cav-1*(+/+), *Rassf1a*+/+;*Cav-1*(-/-) and *Rassf1a*-/-;*Cav-1*(-/-) mice showed alterations in mammary gland architecture such as enlarged, over-sized ductal lumens with epithelial hyperplasia and fibrosis (Figure 1 (A)-(D)), demonstrating that *Rassf1a* and *Cav-1* play important roles in the proper development of the mammary gland. Estrogen and the ERα are of central importance for the formation of the mammary gland, with ERα controlling the growth and elongation of the ductal structures (Bocchinfuso et al., 2000; Bocchinfuso and Korach, 1997; Korach et al., 1996; Mallepell et al., 2006; Sternlicht, 2006). We observed no differences in the number of ER-positive cells between wild-type and *Rassf1a*-/-;*Cav-1*(+/+) mice, and a significant increase in ERα-positive epithelial cells only in the mammary glands of *Cav-1*-deficient *Rassf1a*+/+;*Cav-1*(-/-) and *Rassf1a*-/-;*Cav-1*(-/-) mice (Table 6, Figure 5 (A)-(D)). It is nevertheless conceivable that loss of *Rassf1a* alone or in combination with loss of *Cav-1* could regulate the activity of ERα. RASSF1A inhibits signaling pathways that promote estrogen-independent growth, such as Akt and MAPK, in ER+ breast cancer cells (Roßwag et al., 2020; Thaler et al., 2012). We therefore hypothesized that in the absence of RASSF1A, signaling pathways might be activated that enable estrogen-independent activation of ERα, or enable activation of ERα in the presence of very low estrogen concentrations. To investigate the hypothesis that loss of *Rassf1a* with or without concomitant loss of *Cav-1* leads to activation of the ERα in the absence of estrogen, mammary fat pads were excised from male mice of all four genotypes and examined for the possible presence of mammary ductal structures using whole mounts and H&E-stained paraffin sections (Figure 6 a)-h)) and (Figure 7 (A)-(D)).

**Figure 6.**
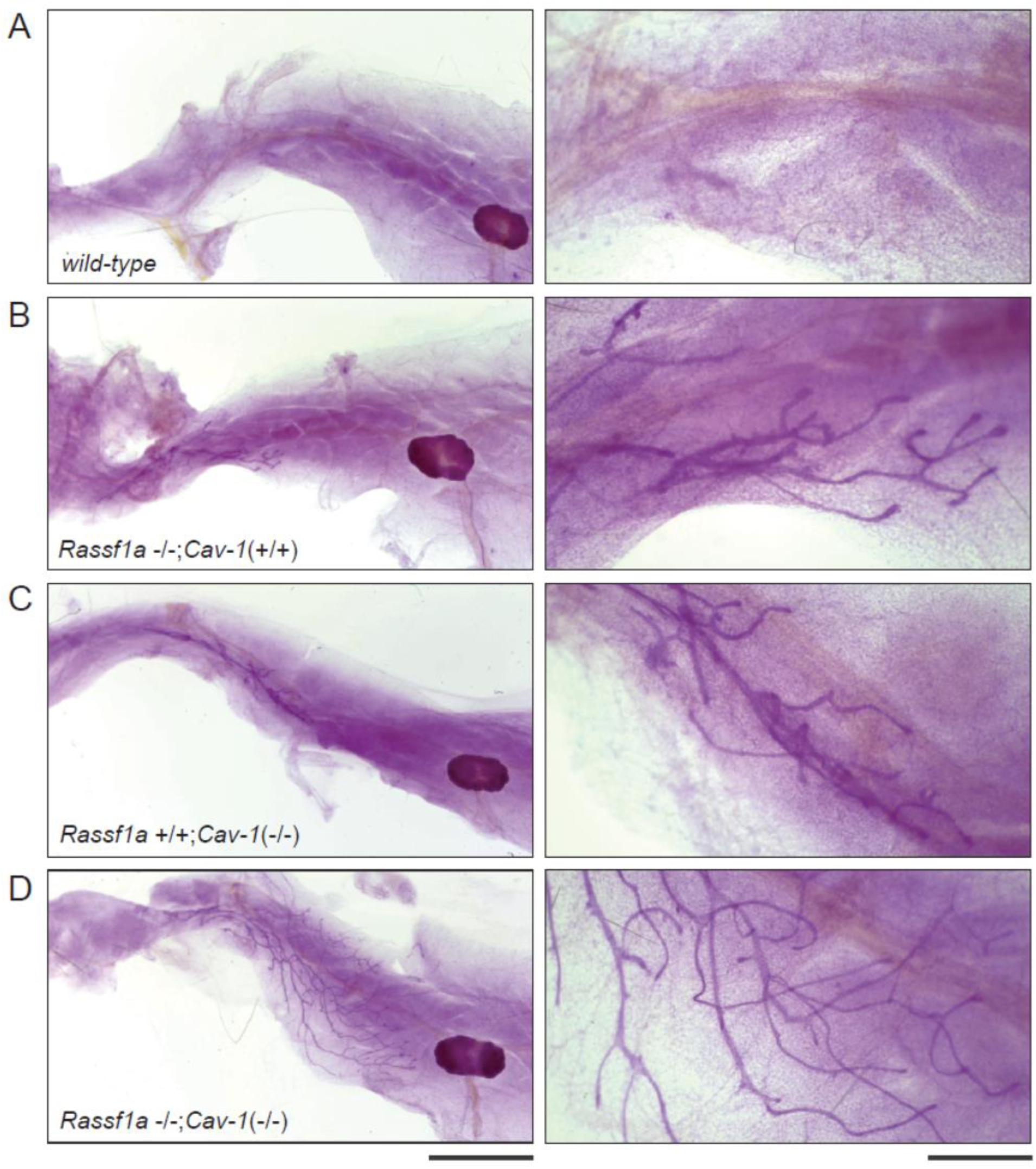
Mammary glands from male *Rassf1a/Cav-1 wild-type*, single knockouts and double knockouts show differences in size and appearances. Whole mount fat pads with mammary gland structures were visualized by staining with carmine alum. (A)-(D) Representative pictures of stained mammary glands from comparable male mice of the same age. (A) *wild-type*, (B) *Rassf1a*-/-;*Cav-1*(+/+), (C) *Rassf1a*+/+;*Cav-1*(-/-), (D) *Rassf1a*-/-;*Cav-1*(-/-). (A)-(D) left panel Bar = 0.5 cm, right panel Bar = 200 µm

**Figure 7.**
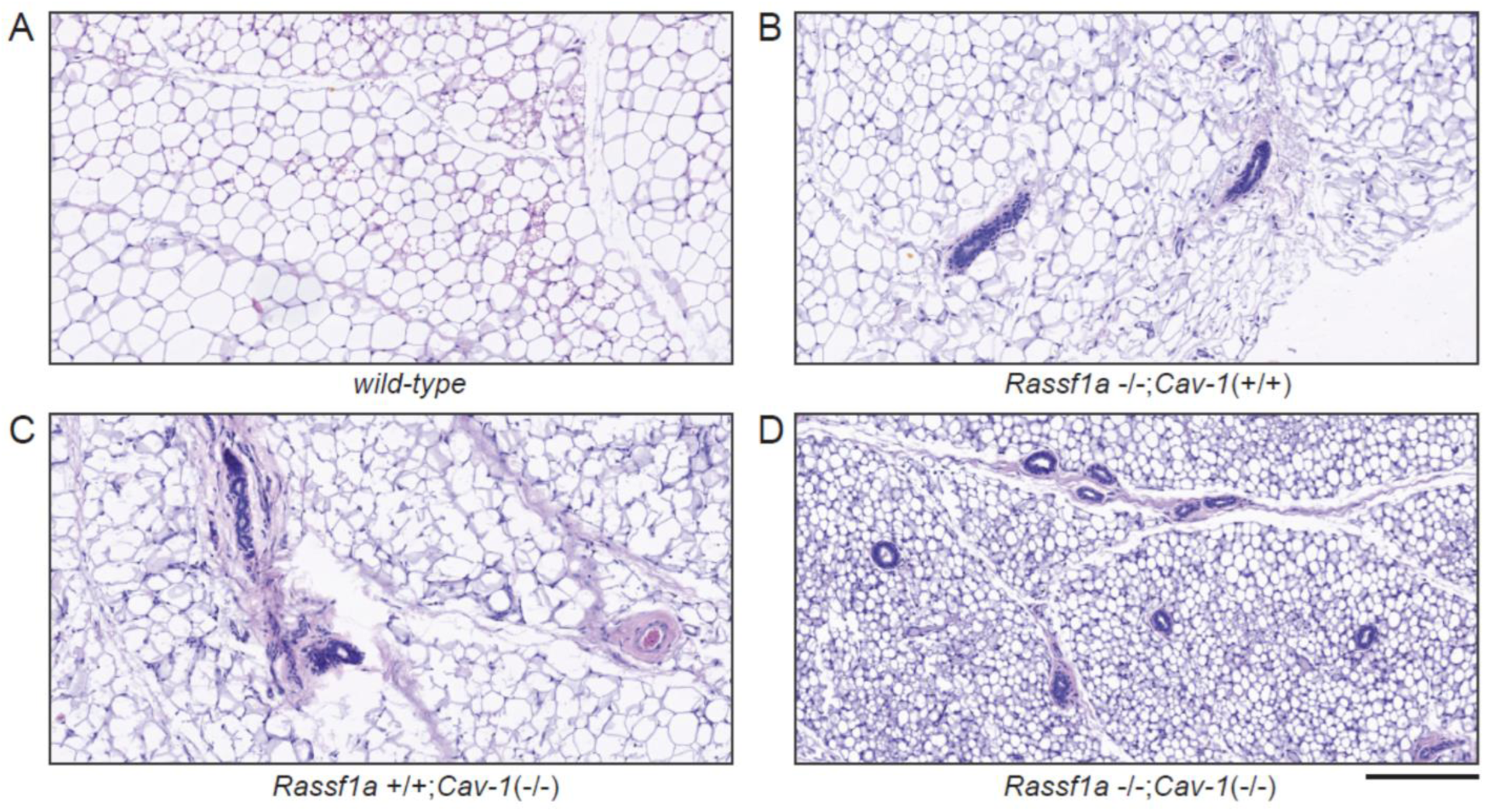
Histopathological analysis of the male mammary fat pads. Representative pictures of H&E-stained sections from paraffin-embedded formalin-fixed fat pads from male mice. Mammary ductal-like structures were only found in the *Rassf1a* and/or *Cav-1*-targeted male mice. (A) *wild-type*, (B) *Rassf1a*-/-;*Cav-1*(+/+), (C) *Rassf1a*+/+;*Cav-1*(-/-), and (D) *Rassf1a*-/-;*Cav-1*(-/-) mice. Bar = 200 µm

Mammary fat pads from a total of 21 male mice were analysed. Interestingly, ductal mammary gland-like structures were only found in the fat pads of *Rassf1a*-/-;*Cav-1*(+/+), *Rassf1a*+/+;*Cav-1*(-/-) and *Rassf1a*-/-;*Cav-1*(-/-) male mice, but not in wild-type males. In the mammary fat pads of *Rassf1a*-/-;*Cav-1*(+/+) males, more mice had developed a rudimentary mammary gland than in *Rassf1a*+/+;*Cav-1*(-/-) males (80% versus 20%) (Table 7).

In *Rassf1a*-/-;*Cav-1*(-/-) males, mammary glands were found in all males analyzed, and in half of these mice, the mammary gland ductal structures had a length of at least one cm. Thus, the *Rassf1a*-/-;*Cav-1*(-/-) male mice have significantly longer ducts than the *Rassf1a*+/+;*Cav-1*(-/-) and *Rassf1a*-/-;*Cav-1*(+/+) males. These observations support the hypothesis that Rassf1a exerts a regulatory function on ERα activity and that loss of *Rassf1a* leads to the activation of ERα in the absence of estrogen or in the presence of very low estrogen concentrations. Since Cav-1 also acts as a regulator of ERα and its loss leads to hypersensitivity to estrogen, the loss of both proteins appears to cause greater activation of ERα than when only one of the two tumor suppressors is missing.

### Teratomas occur in Rassf1a-/-;Cav-1(-/-) female mice

Teratomas are a rare type of tumor that are mostly benign, and which contain immature and fully formed tissues. They arise from pluripotent stem cells or primordial germ cells mostly in the gonads (ovaries or testicles) or along the midline of the body from the remnants of embryonic development. The most common type in females are ovarian teratomas, which are mostly cystic teratomas filled with fluid that contain different types of tissue such as bone, muscle, endocrine glandular tissue and teeth. Although ovarian teratomas can become extremely large, they are benign in the vast majority of cases, and cancerous cases are very rare (Cong et al., 2023).

In this study, ovarian teratomas were found in two out of twelve *Rassf1a*-/-;*Cav-1*(-/-) female mice that were older than 12 months. The teratomas had a size of approximately 1.5 cm in diameter and had a cystic structure, which is typical for ovarian teratomas (Figure 8 (A) and (C)).

**Figure 8.**
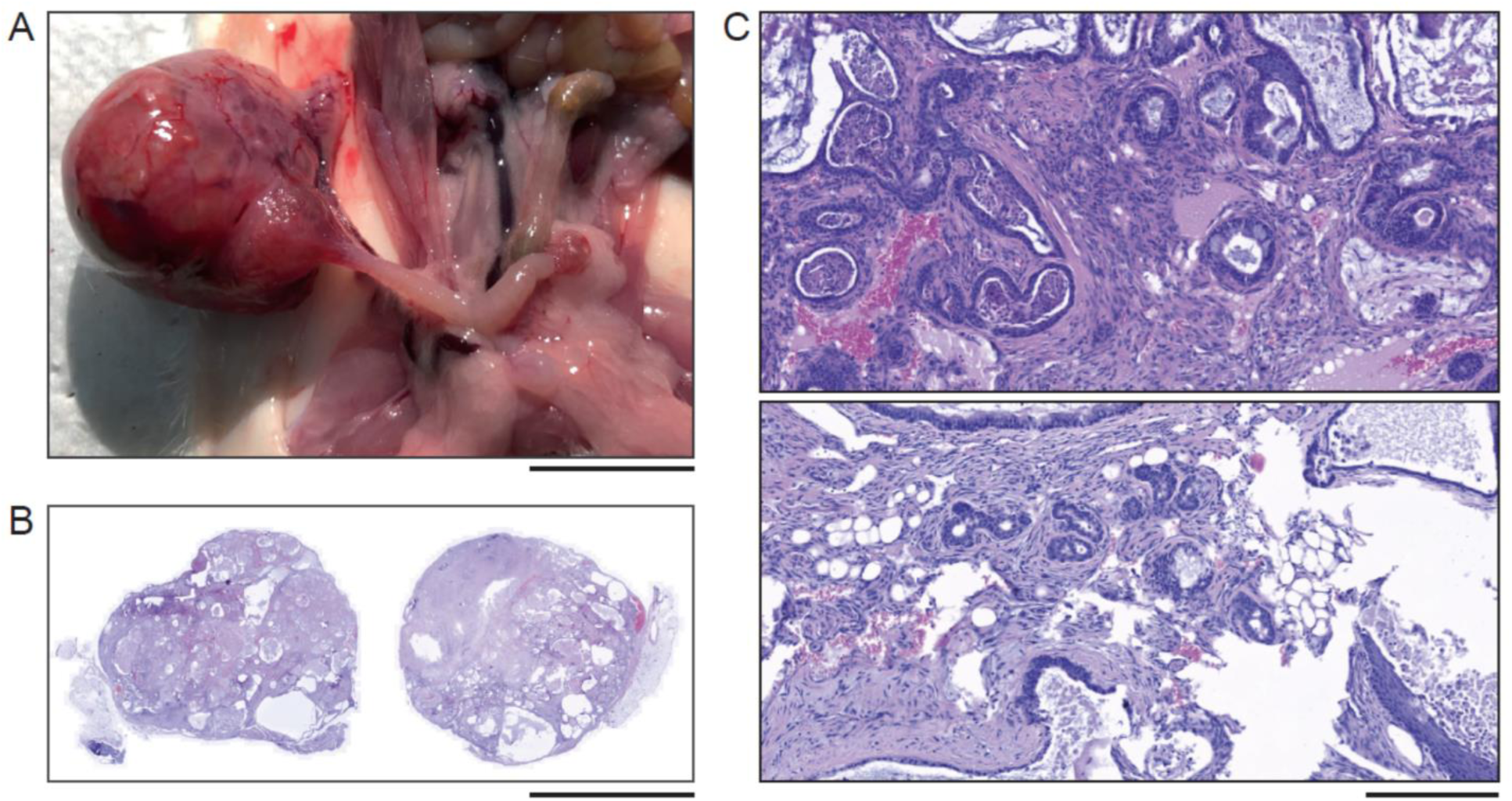
Teratoma from female *Rassf1a-/-/Cav-1(-/-)* mice. (A) Photograph of a teratoma found in *Rassf1a*-/-/*Cav-1*(-/-) mice. Bar = 1 cm. (B) H&E-stained paraffin sections of a teratoma. Both paraffin sections are from different levels within the teratoma. A cystic structure characteristic of ovarian teratomas is visible. Bar = 1 cm. (C) H&E-stained paraffin sections of the teratoma under magnification (upper and lower panel). In addition to many cysts filled with fluid and secretions, many different tissue types and cartalige are visible, such as fat, glandular, squamous, and epithelial tissue. Necrotic tissue and immune cell infiltration are also present. Bar = 200 µm

In addition, these teratomas had many cysts filled with fluid and secretions, and included many different tissue types, such as fat, glandular, squamous, and epithelial tissue. Necrotic areas and immune cell infiltration are also present. However, the teratomas did not show any signs of malignancy and did not include cancerous tissue. These results are consistent with the notion that the concomitant loss of *Rassf1a* and *Cav-1* plays an important role in suppressing benign, premalignant tissue alterations, but that further triggers are required for the malignant transformation of cells and subsequently the development of malignant carcinomas. Likely, the additional activation of oncogenes is required for the development of full-blown invasive carcinomas.

## Discussion

Here we report that genetic loss of the tumor suppressors Rassf1a and Caveolin-1 in transgenic mice leads to distinct, genotype-specific premalignant alterations in mammary gland architecture, including ductal cystic and ductal cystic–acinar hyperplasia, but does not result in invasive breast carcinoma even during pregnancy. Cav-1 deficiency, with or without additional loss of Rassf1a, is associated with increased ERα-positive epithelial cells and hypersensitivity to hormonal cues, whereas Rassf1a loss alone primarily perturbs mammary gland architecture and appears to modulate ERα activity rather than expression. In male mice, loss of Rassf1a and/or Cav-1 permits mammary ductal outgrowth in otherwise rudimentary fat pads, suggesting estrogen-independent or low-estrogen activation of ERα signalling. Collectively, these findings indicate that Rassf1a and Cav-1 are required to maintain normal mammary gland structure and restrain hormone-driven premalignant change, but that additional oncogenic events and/or chronic hormonal stimulation are necessary for progression to full-blown luminal ER+ breast cancer. Intriguingly, development of ovarian teratomas in aged double-knockout females further suggest a broader role for these proteins in germ cell differentiation and meiotic fidelity.

In contrast to previously published *in vivo* studies, we did not expose *Rassf1a*-/- mice to carcinogens that could potentially enhance or accelerate mammary tumor development. In the absence of exposure to carcinogens, loss of *Rassf1a* expression within *Rassf1a-/-*;*Cav-1*(+/+) mice led to ductal cystic hyperplastic (DCH) lesions in the mammary glands (Figure. 1 (B) and Figure. 2 and Table 1), providing support for the notion that RASSF1A suppresses the formation of the pre-malignant precursors of luminal breast carcinogenesis. Similarly, *Cav-1*-deficiency with or without concomitant loss of *Rassf1a* causes DCAH lesions (Figure 2 and Table 1). the additional loss of *Rassf1a* does not lead to significant differences in the appearance of breast lesions between *Rassf1a-/-*;*Cav-1*(+/+) and *Rassf1a-/-*;*Cav-1*(-/-) female mice, and the loss of both tumor suppressors was not sufficient to induce the formation of mammary carcinomas. A possible explanation for this finding might be that both tumor suppressors control similar signaling pathways, and that the progression of pre-malignant lesions requires the loss of additional tumor suppressive mechanisms, such as the loss of DNA repair mechanisms, which leads to the accelerated and increased generation of mutations and subsequent activation of oncogenes. This reflects a common observation that the inactivation of tumor suppressors in in vivo tumor models do not directly lead to the development of invasive carcinoma, and that one or more oncogenes must be activated in parallel for carcinogenesis to occur. It is therefore likely that malignant mammary tumors would have had developed in the *Rassf1a-/-*;*Cav-1*(+/+), the *Rassf1a-/-*;*Cav-1*(+/+) and/or in the *Rassf1a*-/-;*Cav-1*(-/-) mice if they had been either treated with pro-carcinogenic, mutagenic agents, or if they had been exposed to factors such as particular hormones that can promote the development of luminal breast tumors.

Two hormones that are associated with the development of breast cancer are prolactin (Grible et al., 2021; Hathaway et al., 2023; Schuler and O’Leary, 2022; Wang et al., 2016) and progesterone (Coelingh Bennink et al., 2023; Simões et al., 2025). Both hormones play a role during mammary gland development, pregnancy and lactation (Brisken and Scabia, 2020; Schuler and O’Leary, 2022). Progesterone primarily leads to an increased proliferation of stem cells in breast tissue (Brisken and Scabia, 2020). Synthetic progesterone derivatives such as medroxyprogesterone acetate (MDA) are associated with an increased risk of developing breast cancer in humans, and promote the development of breast cancer in mouse experiments if combined with carcinogenic substances (Lanari et al., 2009). Prolactin may similarly foster the development of breast cancer (Schuler and O’Leary, 2022). Prolactin activates JAK2 and STAT5 signaling which is involved in the proliferation, differentiation, and survival of mammary gland epithelial cells (Tian et al., 2020; Xie et al., 2002). Dysregulation of JAK2-STAT5 activity leads to mammary gland developmental defects and/or diseases, including breast cancer (Tian et al., 2020). *Cav-1* plays a central role in regulating the activity of the JAK2/STAT5 signaling pathway (Park et al., 2002). Consistently, we observed that *Cav-1*-deficient, non-pregnant *Rassf1a*+/+;*Cav-1*(-/-) and *Rassf1a*-/-;*Cav-1*(-/-) mice develop DCAH lesions, and that some mice had isolated milk drops in the ducts and ductal-alveolar structures, indicative of non-specific activation of the JAK2/STAT5 signaling pathway and premature milk production (precocious lactation) (Figure 2 and Table 1). Notably, two female *Rassf1a-/-*;*Cav-1*(-/-) mice that were neither pregnant nor lactating exhibited a massive secretory DCAH phenotype with structures that were filled with milk drops (Figure 3 (A)-(D)). This observation is consistent with the notion that loss of both Cav-1 and Rassf1a can coordinately foster precocious milk production in the mammary glands of female *Rassf1a-/-*;*Cav-1*(-/-) mice. However, given the small number of animals that exhibit this histological pattern, further work would be needed to substantiate this notion.

To determine whether the additional loss of *Rassf1a* in *Cav-1*-deficient mice under the influence of progesterone or prolactin is associated with malignant transformation and the development of invasive lesions, we examined the mammary glands from pregnant mice of all four genotypic groups. However, neither *Rassf1a*+/+, *Cav-1*(-/-) nor *Rassf1a*-/-;*Cav-1*(-/-) mice develop breast cancer during pregnancy (Figure 4 (A)-(D)). These results demonstrate that concomitant loss of *Rassf1a* and *Cav-1* is not sufficient to initiate breast cancer development following short-term exposure to progesterone or prolactin during pregnancy, and that further changes, most likely the activation of oncogenes, and/or chronic exposure to prolactin or progesterone are important for the development of mammary carcinomas.

Previous studies reported that *Cav-1*-deficient mice exhibit increased ERα expression in mammary epithelial cells, and that mammary glands from *Cav-1* (-/-) mice are functionally hypersensitive to the effects of estrogen exposure (Mercier et al., 2009). RASSF1A inhibits the expression and function of ERα in breast cancer cells partly via the same signalling pathways as Cav-1 (Thaler et al., 2012), suggesting that loss of *Rassf1a* and *Cav-1* should lead to increased expression of ERα in *Rassf1a*- and *Cav-1*-deficient mice. However, a significant increase in ERα expression and an increase in ER+ epithelial cells in the mammary glands was only observed in *Rassf1a*+/+,*Cav-1*(-/-) and *Rassf1a*-/-;*Cav-1*(-/-) mice (Figure 4 (A)-(D)) and Table 6).

Although no differences in the number of ERα-positive cells were observed when mammary glands from wild-type and *Rassf1a*-/-;*Cav-1*(+/+) mice were compared, and a significant increase in ERα+ epithelial cells was only found in the mammary glands of *Cav-1*-deficient mice (Table 6, Figure 5 (A)-(D)), it is nevertheless conceivable that loss of *Rassf1a* alone or in combination with loss of *Cav-1* could have an influence on the activity of ERα. In human ER+ breast cancer cells, RASSF1A inhibits signaling pathways that promote estrogen-independent growth, such as AKT and MAPK (Roßwag et al., 2020; Thaler et al., 2012). Thus, in the absence of RASSF1A, signaling pathways may be activated that enable estrogen-independent activation of ERα, or enable activation of ERα in the presence of very low estrogen concentrations. Consistent with this notion, ductal mammary gland-like structures in the fat pads of male mice were only found in *Rassf1a*-/-;*Cav-1*(+/+), *Rassf1a*+/+;*Cav-1*(-/-) and *Rassf1a*-/-;*Cav-1*(-/-) mice, but not in wild-type males (Figure 6 (A)-(D)) and (Figure 7 (A)-(D) and Table 7). As estrogen and the ERα are of central importance for the formation of the mammary gland, with ERα controlling the growth and elongation of the ductal structures (Bocchinfuso et al., 2000; Bocchinfuso and Korach, 1997; Korach et al., 1996; Mallepell et al., 2006; Sternlicht, 2006), these observations support the hypothesis that loss of Rassf1a leads to the activation of ERα in the absence of estrogen or in the presence of very low estrogen concentrations. Since Cav-1 also acts as a regulator of ERα and its loss leads to hypersensitivity to estrogen (Mercier et al., 2009), the loss of both proteins appears to cause greater activation of ERα than if only one of the two tumor suppressors is missing.

A limitation of this study is that the phase of the estrous cycle in which the mice were at the time of mammary gland removal was not determined. The estrus cycle in mice is a 4 - 5 days reproductive cycle consisting out of four different stages referred to as proestrus, estrus, metestrus and diestrus. The estrus cycle is characterized by rapid changes in estradiol and progesterone levels, accompanied by changes in the expression levels of ERα and progesterone receptor (Silberstein et al., 2006). A possible explanation for the equal distribution of the wild-type and *Rassf1a*-/-;*Cav-1*(+/+) mice between the weak, moderate, and strong ERα groups could therefore be that the mice were in different phases of the estrous cycle and therefore differed in ERα expression. Since the *Rassf1a*-/-;*Cav-1*(+/+) mice also show differences in ERα expression between individual mice, it is likely that loss of Rassf1a has no influence on the fluctuation of ERα expression during the estrous cycle.

Mature cystic ovarian teratomas without any signs of malignancy were found in two out of twelve *Rassf1a*-/-;*Cav-1*(-/-) female mice aged more than 12 months (Figure 8 (A)-(C)). Teratomas are tumors that arise from pluripotent stem cells or primordial germ cells. *Rassf1a* and *Cav-1* play important roles in stem cell differentiation (Baker and Tuan, 2013; Dalton et al., 2023; Papaspyropoulos et al., 2018). It can therefore be speculated that the occurrence of ovarian teratomas in the group of Rassf1a-/-;Cav-1(-/-) females could possibly be explained by the fact that concomitant loss of *Rassf1a* and *Cav-1* leads to a larger number of primordial germ cells that do not differentiate, thereby increasing the likelihood that one of these cells will later develop into a teratoma. Mature cystic teratomas derive from primordial germ cells that undergo parthenogenic activation. These germ cells typically complete the first meiotic division but fail in subsequent divisions, leading to tumors with a 46,XX karyotype (Surti et al., 1990). Thus, mechanisms that foster failure of proper meiotic cell division are essential for ovarian teratoma development. RASSF1A coordinates and regulates the anaphase progression, mainly by controlling the activity of the anaphase-promoting complex/cyclosome (APC/C) (Song et al., 2004). In addition, RASSF1A plays a critical role in the kinetochore-spindle complex, particularly during mitosis and meiosis. RASSF1A is a microtubule-associated protein, which maintains microtubule stabilization, spindle organization and attachment of microtubules to the kinetochores, thereby ensuring proper mitotic progression (Jeon and Oh, 2020; Liu et al., 2003; Song et al., 2004). RASSF1A depletion causes premature APC/C activation and microtubule instability, which results in accelerated mitotic progression and enhanced risk of failure in chromosome separation, leading to mitotic abnormalities and aneuploidy. Therefore, *Rassf1a*-depletion might increase the risk of failure of meiotic divisions. Since teratomas were only observed in *Rassf1a-/-*;*Cav-1*(-/-) females, it is likely that *Cav-1* depletion additionally increases the risk of mitotic division failure. This notion is supported by studies showing that Cav-1 is involved in meiotic cell cycle progression (Sato et al., 2006). However, further research such as karyotype analysis would be required to demonstrate a functional correlation between the loss of Rassf1a and Cav-1, defective meiotic divisions, and an increased incidence of teratomas.

In summary, our data show that genetic loss of the tumor suppressors Rassf1a and Caveolin-1 in mice causes distinct premalignant changes in mammary gland architecture, but is insufficient on its own to drive invasive breast cancer. Cav-1 loss increases the number of ERα-positive epithelial cells and hormonal hypersensitivity, while Rassf1a loss mainly disrupts gland structure and modulates ERα activity. Accordingly, their loss in male mice enables mammary ductal outgrowth even under low-estrogen conditions. Together, the data suggest that Rassf1a and Cav-1 help maintain normal mammary structure and limit hormone-driven premalignant changes, that additional oncogenic events or chronic hormonal exposure are required for luminal ER+ tumor progression, and that these proteins may also have roles in germ cell differentiation and meiotic fidelity.

### Limitations of the study

This study demonstrates that Rassf1a and Cav-1 contribute to maintaining the normal structure of the mammary gland and limiting hormone-induced premalignant changes in our mouse models. However, a limitation of this study is that we cannot determine which additional factors, aside from the loss of Rassf1a, are necessary for the development of full-blown ER+ breast carcinomas. The additional loss of Cav-1, which also inhibits ERα activity, did not result in fully developed breast carcinomas in our mouse models either. A possible explanation for this could be that Rassf1a and Cav-1 regulate similar signaling pathways, and the inactivation of both tumor suppressors has mainly additive effects. We hypothesize that the activation of oncogenes is necessary for the development of full-blown ER+ breast carcinomas. Key oncogenes in luminal breast carcinogenesis include activating mutations in components of the PI3K/Akt signaling pathway. Since Rassf1a inhibits the PI3K/Akt signaling pathway in *in vitro* experiments, it is plausible that Rassf1a could also have an inhibitory effect on the PI3K/Akt signaling pathway even when it harbors activating mutations, thereby inhibiting the carcinogenesis of luminal breast cancers. However, in this study we did not investigate whether activating mutations in the PI3K/Akt signaling pathway in Rassf1a-/- mice, compared to Rassf1a wild-type mice, lead more frequently or more rapidly to the development of luminal breast carcinomas in Rassf1a-/- mice compared to Rassf1a wild-type mice. Thus, we can only speculate on this at this time and will focus on this question in the future.

Another limitation of this study is the small number of *Rassf1a*-/-;*Cav-1*(-/-) females that were older than 12 months, and the fact that two of the 12 females analyzed had teratomas. Although Rassf1a and Cav-1 regulate cellular processes that are important for proper meiotic cell cycle progression, and it is therefore plausible that the loss of both proteins could promote the development of teratomas, further research will be required to demonstrate a functional correlation between the loss of Rassf1a and Cav-1, defective meiotic divisions, and an increased incidence of teratomas.

## STAR*METHODS

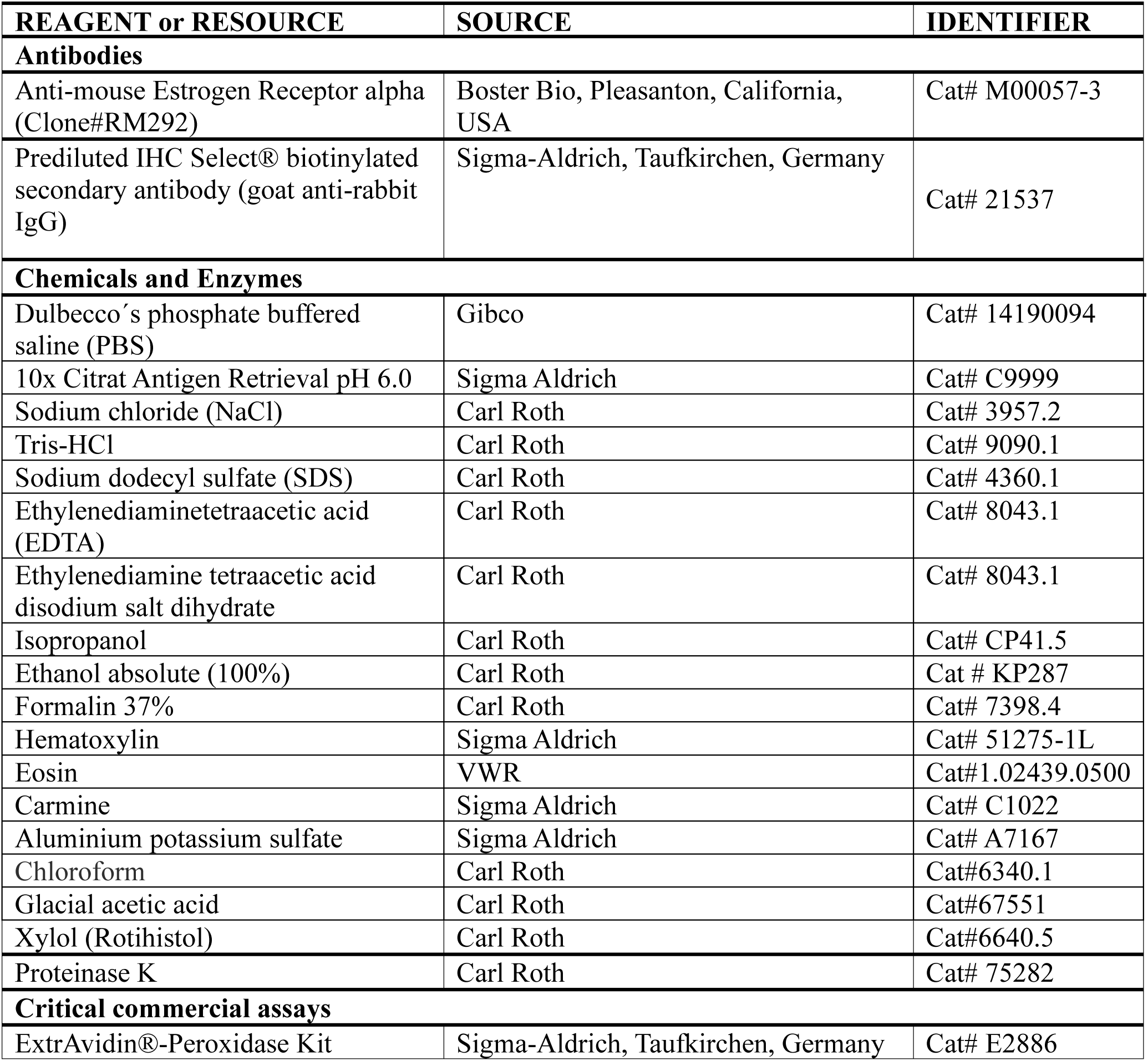
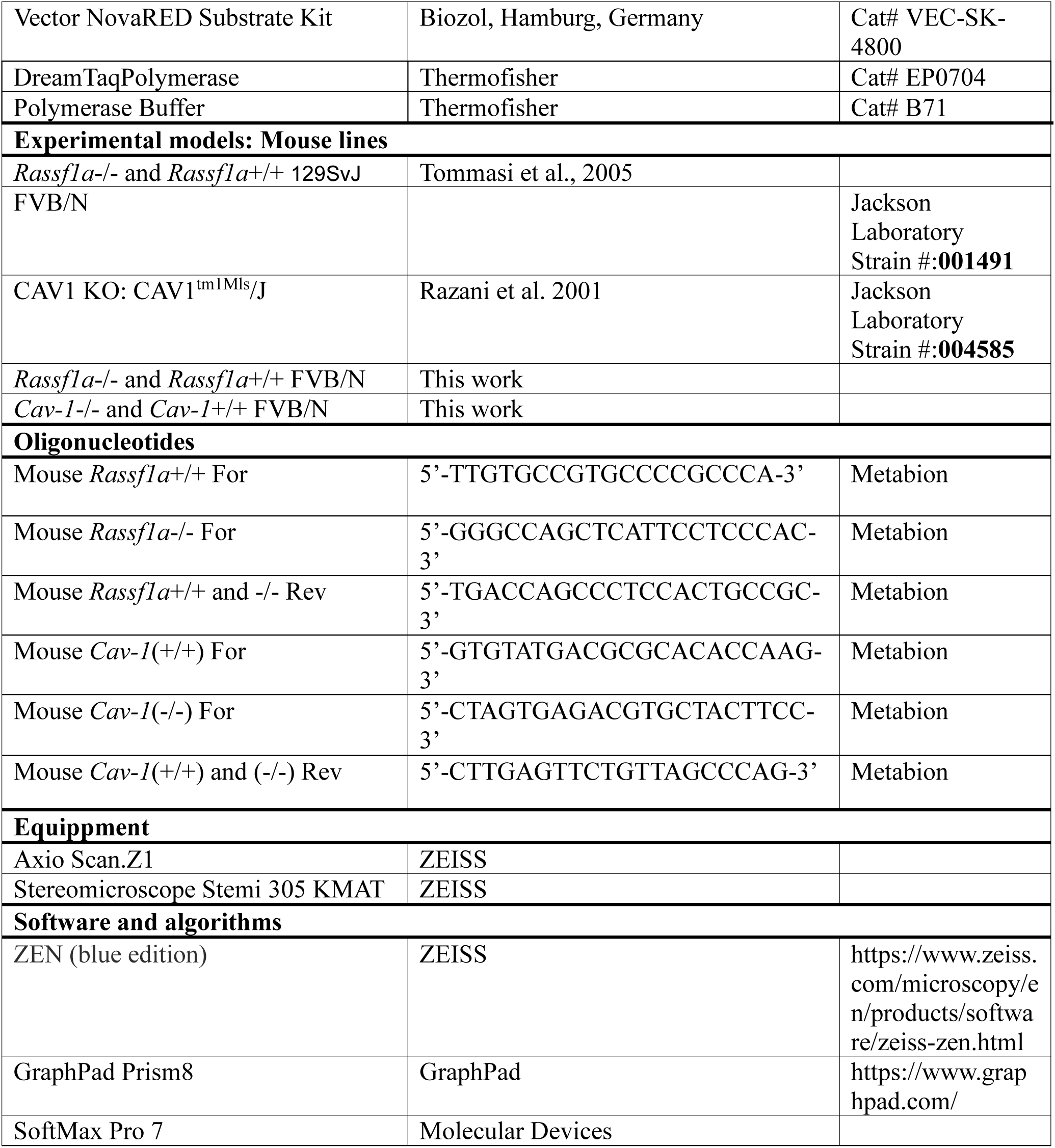

## RESOURCE AVAILABILITY

### Lead contact

Further information and requests for resources and reagents should be directed to and will be fulfilled by the Lead Contact, Sonja Thaler.

## Material availability

Material used in this study will be available for academic researchers upon completion of the Material Transfer Agreement.

## Data and code availability

Any information required to reanalyze the reported data is available from the lead contact upon request.

## EXPERIMENTAL MODEL AND SUBJECT DETAILS

### Mouse mammary tumor models

*Rassf1a*-/- and *Rassf1a*+/+ 129SvJ mice (Tommasi et al., 2005) were provided by Gerd Pfeifer from Van Andel Institute, Michigan, USA, and back-crossed onto the FVB strain for at least 6-10 generations. Heterogenous *Rassf1a*+/- mice were intercrossed to obtain homozygous *Rassf1a*-/- and wild-type *Rassf1a*+/+ littermates. *Cav*-1-deficient (CAV1 KO: CAV1^tm1Mls^/J) mice (Razani et al., 2001) purchased from Jackson Laboratory (Bar Harbor, Maine, USA) were backcrossed onto the FVB strain for more than 12 generations, and are referred to as *Cav*-1(-/-) mice. To generate *Rassf1a*+/+; *Cav-*1(-/-) and *Rassf1a*-/-;*Cav-*1(-/-) mice, *Cav-*1(-/-) animals were crossed with homozygous *Rassf1a*-/- and wild-type *Rassf1a*+/+ mice.

### Genotyping of transgenic mice

PCRs for genotyping were performed with genomic DNA isolated from the tips of the tails of the mice. Therefore, a piece of mouse tail approximately 1-2 mm long is placed in a sterile Eppendorf tube and mixed with 400 µl of lysis buffer (20 mM Tris-HCl, pH 8.0, 400 mM NaCl, 5 mM EDTA, 1% SDS) and proteinase K (final concentration 0.25 mg/ml). The tubes are incubated overnight at 55 °C with shaking (750 rpm), centrifuged afterwards (13,000 rpm, 10 min) and 350 µL of the supernatant was transferred into a new tube. To precipitate the genomic DNA 250 µl of isopropanol was added, incubated for 10 min and centrifuged (13,000 rpm, 15 min). The supernatant is discarded, the DNA pellet then air-dried for 1–2 min and resuspended in 100 µl H_2_O. PCR was performed by using the following primer pairs: *Rassf1a*+/+ fw: 5’-TTGTGCCGTGCCCCGCCCA-3’, *Rassf1a*-/- fw: 5’-GGGCCAGCTCATTCCTCCCAC-3’, *Rassf1a* rev: 5’-TGACCAGCCCTCCACTGCCGC-3’, *Cav-1*(+/+) fw: 5’-GTGTATGACGCGCACACCAAG-3’, *Cav-1*(-/-) fw: 5’-CTAGTGAGACGTGCTACTTCC-3’, *Cav-1* rev: 5’-CTTGAGTTCTGTTAGCCCAG-3’. All mice were housed and maintained in a pathogen-free environment following the German national guidelines of the animal protection law.

### Histological analysis

Mammary glands were excised, fixed in 4% buffered formalin for 18-24 hours, dehydrated and embedded in paraffin. Sections with a thickness of 4-6 µm were re-hydrated and antigen retrieval was performed using citrate buffer pH 6.0. Formalin-fixed and paraffin-embedded mammary glands were used for hematoxylin and eosin (H&E) or immunohistochemical staining. Mouse ERα expression was detected using the mouse-specific anti-ER alpha ESR1 Rabbit Monoclonal Antibody, Clone#RM292 (Catalog # M00057-3) (Boster Bio, Pleasanton, California, USA) (1:200 dilution in IHC/ICC Blocking Reagent High Protein, Thermo Fisher Scientific GmbH, Dreieich, Germany. Staining was visualized using the prediluted IHC Select® biotinylated secondary antibody (goat anti-rabbit IgG), Sigma-Aldrich, Taufkirchen, Germany and the avidin-biotin complex (ABC) method by using the ExtrAvidin®-Peroxidase (1:100 diluted), Sigma-Aldrich, Taufkirchen, Germany, with NovaRED (Biozol, Hamburg, Germany) as the chromogen. To ensure that the immunohistochemical staining for ERα expression on the paraffin sections of mouse mammary glands was specific and that the staining intensities could be compared, a piece of an ER-negative and an ER-positive human breast carcinoma was additionally applied to each individual slide and also stained. ERα positive cells were quantified on the basis of percentage positivity of luminal cells on each section. For ERα expression only nuclear reactivity was taken into account. Imaging of stained sections was performed by using the ZEISS Axio Scan.Z1 and ZEN (blue edition) software, ZEISS, Jena, Germany.

### Mammary whole mount staining

Inguinal mammary glands (#4 and #5) were excised, spread onto glass slides, and fixed in Carnoy’s fixative composed out of 100% EtOH, chloroform and glacial acetic acid (ratio 6:3:1) for 18-24 h at room temperature. The fixed mammary glands were then washed in 70% EtOH for 15 min and changed gradually to distilled water. Once hydrated, the mammary glands were stained overnight in carmine alum (1 g carmine and 2.5 g aluminium potassium sulfate in 500 ml distilled water). Carimine alum (C1022) and aluminium potassium sulfate (A7167) were purchased from Sigma Aldrich (Taufkirchen, Germany). After carmine alum staining mammary glands were cleared in xylol as previously described (Stanko and Fenton, 2017). Imaging of mammary whole mounts was performed by using ZEISS Stereomicroscope Stemi 305 KMAT or ZEISS Axio Scan.Z1 and ZEN (blue edition) software, ZEISS, Jena, Germany.

### Statistical analysis

Fisher’s exact test was performed using the Microsoft Excel 2010 software package and GraphPad Prism 8.3.1 software. Statistical significance was defined as a p-value of <0.05.

## Acknowledgements

This project was funded by the Deutsche Forschungsgemeinschaft (DFG grant TH1523/3-1 and TH1523/3-3 to S. Thaler) and by the Medical Faculty Mannheim. This research was conducted with the help of the LIMA Core Facility of the Medical Faculty Mannheim.

## Author’s Contributions

Conceptualization: S.T., J.P.S., Data curation and analysis, C.L.C., S.T., S.R., H.T.W., T.W., I.S., Resources, S.T., J.P.S., Supervision, S.T., Founding acquisition, S.T., Writing-original draft, S.T., Writing-review and editing, J.P.S., C.L.C., S.T., S.R., H.T.W., T.W., I.S.

## Ethics declarations

Not required

